# Pan-cancer analysis of the ion permeome reveals functional regulators of glioblastoma aggression

**DOI:** 10.1101/2023.04.07.536030

**Authors:** Alexander T. Bahcheli, Hyun-Kee Min, Masroor Bayati, Weifan Dong, Alexander Fortuna, Hongyu Zhao, Irakli Dzneladze, Jade Chan, Xin Chen, Kissy Guevara-Hoyer, Peter B. Dirks, Xi Huang, Jüri Reimand

## Abstract

Ion channels, transporters, and other ion-permeating proteins, collectively comprising the ion permeome (IP), are common drug targets. However, their roles in cancer are understudied. Our integrative pan-cancer analysis shows that IP genes display highly-elevated expression patterns in subsets of cancer samples significantly more often than expected transcriptome-wide. To enable target identification, we identified 410 survival-associated IP genes in 29 cancer types using a machine learning approach. Notably, *GJB2* and *SCN9A* show prominent expression in neoplastic cells and associate with poor prognosis in glioblastoma (GBM), the most common and aggressive brain cancer. *GJB2* or *SCN9A* knockdown in patient-derived GBM cells induces transcriptome-wide changes involving neural projection and proliferation pathways, impairs cell viability and tumor sphere formation, mitigates tunneling nanotube formation, and extends the survival of GBM-bearing mice. Thus, aberrant activation of IP genes appears as a pan-cancer feature of tumor heterogeneity that can be exploited for mechanistic insights and therapy development.

## INTRODUCTION

Developing rational cancer therapies is a challenge as broad-spectrum therapies fail to target tumor heterogeneity and multiple avenues of cancer progression (1). Molecular profiling as standard of care has advanced precision treatment regimens for many cancer types (2). Despite these advances, developing new therapies remains a long and uncertain process. Conversely, repurposing approved drugs is an appealing alternative with many successes (3), such as the use of the type II diabetes drug metformin as an anti-cancer agent (4).

Ion channels permeate ions across membranes based on ionic electrochemical gradients. Voltage-gated ion channels are regulated by changes in transmembrane voltage potential and are involved in a variety of physiological processes such as neuronal signal transmission and epithelial cell secretion. Ligand-gated ion channels are regulated by chemical messengers such as neurotransmitters at neural synapses and neuro-muscular junctions. Ion transporters actively move ions across membranes through energy consumption and conformational change. Gap junctions create intercellular connections to enable the passage of ions and small molecules between different cells. Collectively, we refer to these proteins as the *ion permeome* (IP). The ion permeome is extensively studied in the context of human disease and are well-recognized drug targets. For example, a common therapy for renal hypertension and cardiovascular disease involves Ca^2+^ ion channel blockers (5,6). IP inhibitors are frequently used as local anaesthetics, such as lidocaine and carbamazepine (7). We and others have uncovered the multifaceted roles of ion channels in regulating tumor cell-intrinsic properties and tumor cell-microenvironment interactions, thereby establishing specific ion channels as therapeutic targets in brain cancer (8–18). Despite the identification of specific ion channels as regulators of malignancy of individual cancer types, a comprehensive interrogation of the transcriptomic landscape and clinical significance of the IP in human cancer has not been achieved.

Glioblastoma (GBM) is the most common and deadliest form of primary brain cancer. Despite multi-modal therapy combining surgery, radiotherapy, and chemotherapy using the DNA alkylating agent temozolomide, median patient survival is only ~15 months (19). GBM is characterised by genetic, molecular, and phenotypic heterogeneity at inter- and intra-tumoral levels. GBM comprises distinct molecular subtypes (mesenchymal, proneural, classical), each with specific genomic mutations, gene expression signatures, and clinical characteristics (20–22). Individual GBM tumors harbor diverse tumor and stromal cell populations. This tremendous degree of tumor heterogeneity drives therapy resistance and tumor recurrence (23,24). As such, there is an urgent need to identify actionable therapeutic targets and treatment opportunities.

Here we analysed the transcriptomic landscape and clinical associations of IP genes across 10,000 human cancer samples. We discovered that IP genes were excessively upregulated in subsets of tumors significantly more than expected, revealing a novel aspect of tumor heterogeneity. Using machine learning, we established a catalogue of IP genes whose elevated expression is associated with patient survival outcomes. In GBM, we focused on two IP genes, *GJB2* and *SCN9A*, and demonstrated their roles in promoting GBM aggression using patient-derived tumor cells and xenograft models. Our study highlights alterations in the IP as a cancer hallmark and provides a useful resource for functional studies of IP genes for therapeutic and biomarker development.

## RESULTS

### Transcriptomic landscape of the ion permeome in cancer

To interrogate IP in cancer, we analysed 9352 cancer transcriptomes of 33 cancer types from the Cancer Genome Atlas (TCGA) PanCanAtlas project (25) (**Table S1**). We studied 276 high-confidence druggable IP genes from the Guide to Pharmacology database (26) (**Figure S1**, **Table S2**). We first investigated pan-cancer expression of IP genes using dimensionality reduction and clustering, which revealed tissue- and disease-type specific patterns (**Figure 1a**). For example, GBM and low-grade glioma (LGG) clustered together as did subtypes of kidney cancers (renal cell carcinoma (KIRC), renal papillary cell carcinoma (KIRP) and kidney chromophobe (KICH)). Organ-specific clustering of other cancer samples by IP genes was also detected. For example, digestive tract-related cancers clustered together such as colorectal, stomach and pancreatic cancers (colon adenocarcinoma (COAD), rectum adenocarcinoma (READ), pancreatic adenocarcinoma (PAAD), stomach adenocarcinoma (STAD)), while several organ systems showed distinct clusters, such as two subtypes of melanoma (skin cutaneous melanoma (SKCM), uveal melanoma (UVM)).

**Figure 1.**
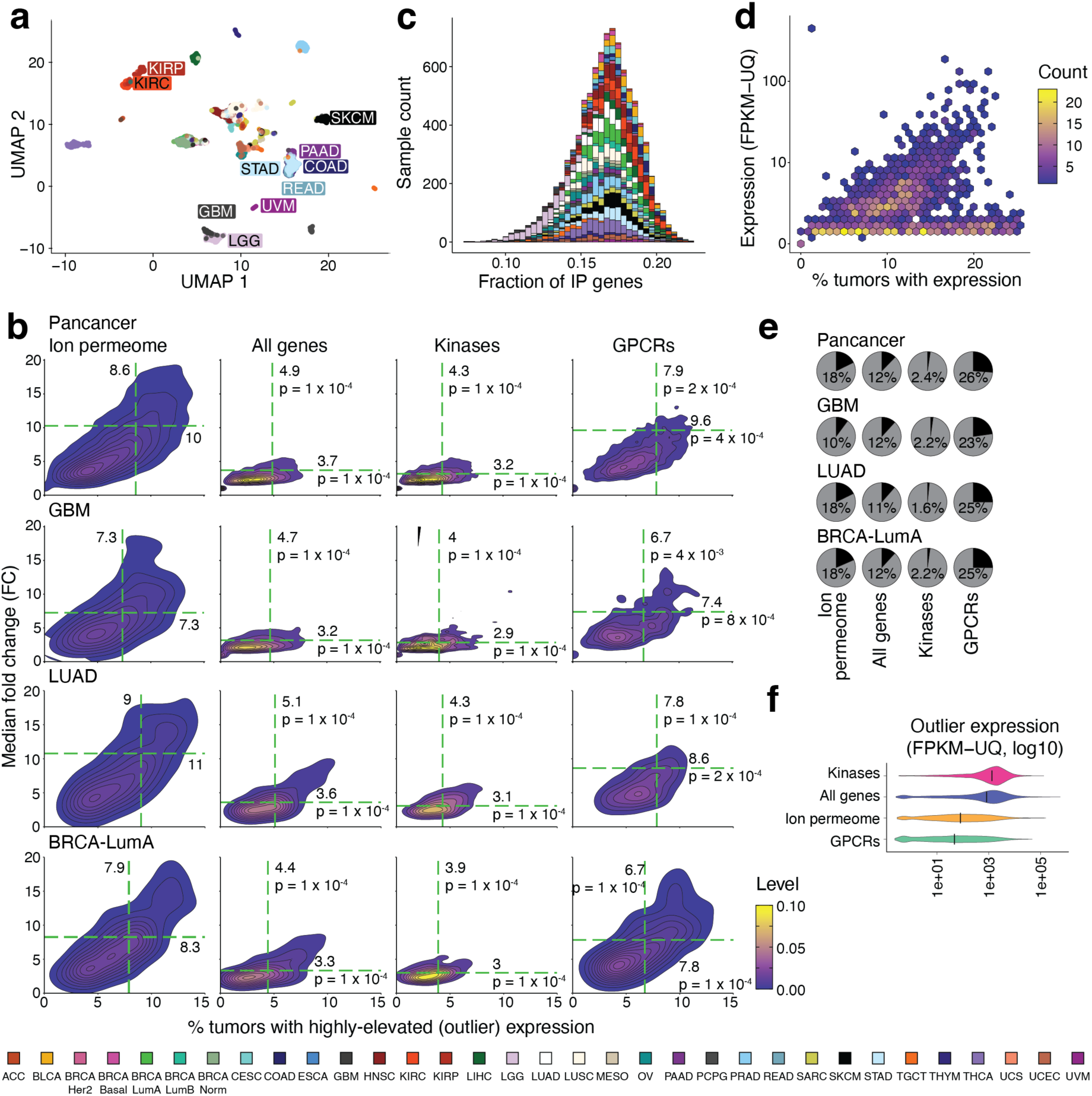
Highly elevated expression of ion permeome genes in cancer. **(a)** Dimensionality reduction analysis of cancer transcriptomes using the expression profiles of ion permeome (IP) genes shows their tissue-specific clustering in cancer. A UMAP projection of 276 IP genes in 33 cancer types from TCGA is shown. **(b)** IP genes are highly upregulated in subsets of cancer samples. Two-dimensional density plots show the joint distribution of gene expression increase (fold-change (FC), log2) and the fraction of cancer samples affected. IP genes (left) are compared to three control analyses (middle to right): (i) all protein-coding genes, and two classes of drug targets: (ii) kinases and (iii) GPCRs. Dashed green lines show median values. Control gene sets were down-sampled to IP gene counts and the representative iterations with median fold-change are shown. **(c)** Histogram of IP genes with highly-elevated expression in each cancer type. Fraction of IP genes with outlier expression for each cancer sample is shown. **(d)** Switch-like expression patterns of IP genes across all cancer types. The 2D density plot shows the fraction of cancer samples with non-zero IP gene expression and the corresponding median non-zero expression values for each gene. **(e)** Pie charts show the fraction of genes with switch-like expression among IPs and control gene sets (all genes, kinases, GPCRs). **(f)** Expression levels of IPs in the highly-elevated groups of samples compared to control gene sets. Control gene sets were down-sampled to IP gene counts and representative iterations with median raw expression are shown.

Transcriptomic analysis revealed dramatic patterns of IP gene overexpression in individual cancer samples. First, we considered IP genes that were expressed in most samples of a given cancer type. A typical IP gene showed 10-fold upregulation in a subset of samples (9%) compared to other samples of the same cancer type (**Figure 1b**). This affected 49 (18%) IP genes per cancer sample on average, based on non-parametric Tukey’s outlier analysis (27) (**Figure 1c**). Overexpression of IP genes was identified in most cancer types in our dataset, as well as pooled pan-cancer dataset. Next, we considered the subset of IP genes with switch-like activation, which showed prominently elevated expression in a minority of cancer samples and no expression in other samples. Switch-like activation affected an additional 18% of IP genes on average (**Figure 1d-e**, **Figure S2a**). A typical IP gene was expressed in hundreds of copies in high-outlier group of cancer samples (median 290 FPKM-UQ), while some genes exceeded these levels by several orders of magnitude (10^4^ – 10^5^ FPKM-UQ) (**Figure 1f**).

To evaluate the significance of IP overexpression in cancer, we repeated the outlier analysis by re-sampling protein-coding genes as controls. In all cancer types, overexpression of IP genes was significantly more pronounced compared to all protein-coding genes, with a median fold-change of 3.7 and 5% of samples affected on average (*P* < 10^−6^, permutation test) (**Figure 1b**, **Figure S2b**). We repeated the outlier analysis using two other major drug target classes: kinases and G protein-coupled receptors (GPCRs). Aberrant overexpression of IP genes significantly exceeded the overexpression of genes encoding kinases. Interestingly, genes encoding GPCRs were highly upregulated in most cancer types, while the extent and frequency of upregulation among IP genes was often significantly higher (**Figure S2b**). Collectively, these data demonstrate that IP genes undergo dramatic upregulation in a fraction of cancer samples, implicating their contributions to tumor heterogeneity and disease mechanisms.

### Survival associations of ion permeome genes in multiple cancer types

To investigate the IP in cancer pathology, we systematically prioritised IP genes that significantly associated with overall or progression-free patient survival in individual cancer types, using a machine learning approach from our previous study (28). Briefly, we trained a collection of Cox proportional-hazards (CoxPH) survival regression models on subsets of cancer samples with IP genes as features, followed by regularisation that selected the most informative IP genes in each model. We nominated recurrently selected IP genes from the model feature sets as our top candidates. Cancer types in TCGA were analysed separately to identify IP genes as disease-specific candidates. We benchmarked the analysis by randomly shuffling patient survival data. As expected, this control experiment revealed significantly fewer and attenuated associations with IP genes, suggesting that our computational framework is appropriately calibrated (**Figure S4**).

Our analysis identified 206 IP genes with 410 associations with patient survival in multiple cancer types (**Figure 2a**). We found 12 IP genes per cancer type on average, while most prognostic associations with IP genes were found in only one or two cancer types (74%) (**Table S3**), which is consistent with tissue-specific clustering of IP expression in cancer (**Figure 1a**). The largest numbers of survival associations were found in prostate cancer and luminal-A subtype of breast cancer (23 and 24, respectively). Elevated IP gene expression associated with worse patient prognosis in most IP genes that we identified (240/410 or 59%). Several top-ranking IP genes were found in multiple cancer types, including *ACCN2, GRIN2D*, and *TRPV3* that associated with poor prognosis in six cancer types, and *P2RX6* with seven cancer types (**Figure S5a**). *P2RX6* encodes a P2X receptor that increases renal cancer cell migration and invasion (29). *ACCN2* has been shown to promote tumor growth and metastasis in breast cancer (30), and *GRIN2D* is an angiogenic tumor marker in colorectal cancer (31). *TRPV3* encodes a transient receptor potential cation-selective channel involved in temperature regulation pathways (32). Collectively, the catalogue of prognostic associations of the IP offers a useful resource for functional studies and biomarker discovery.

**Figure 2.**
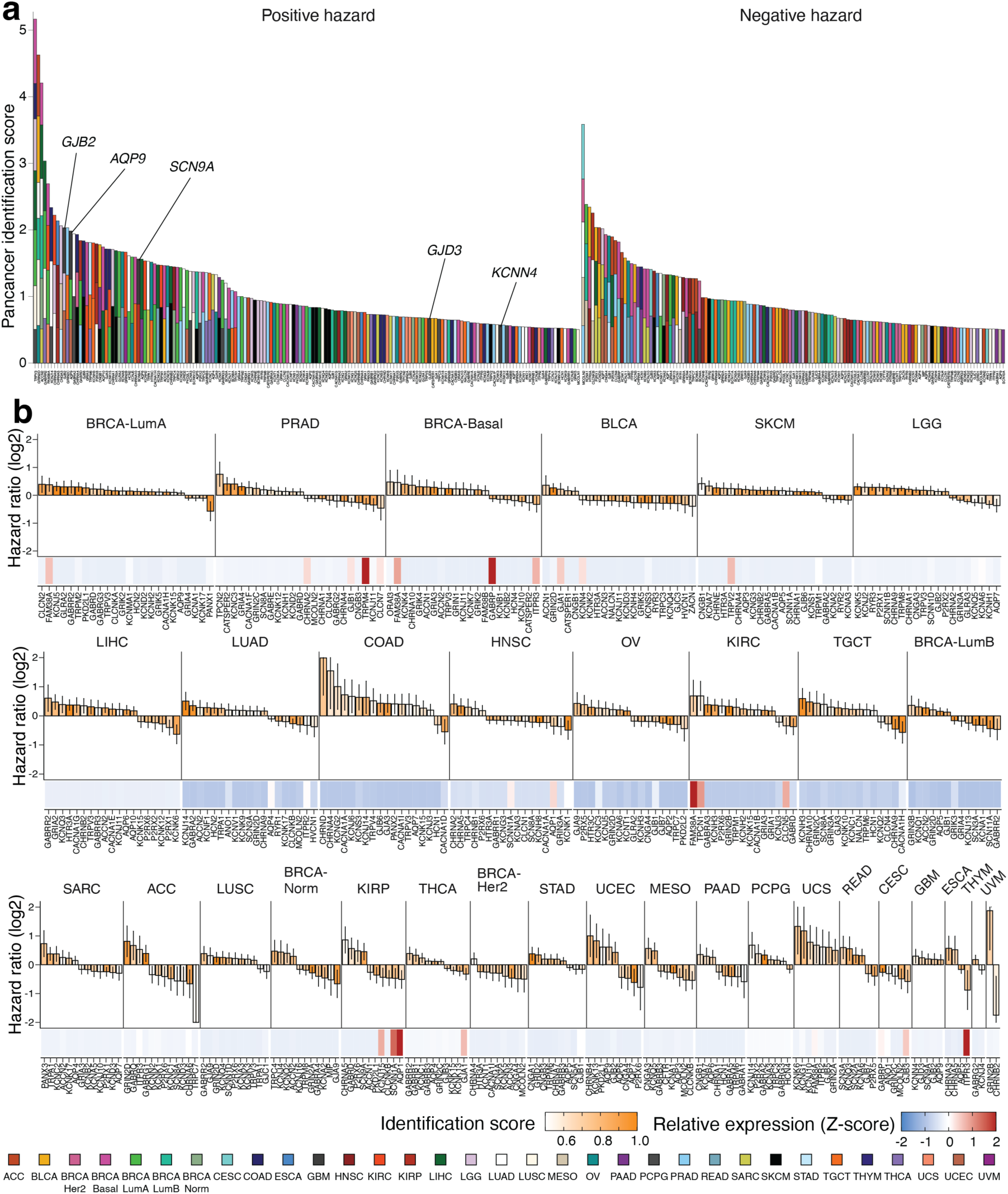
Patient survival associations of ion permeome (IP) genes in cancer. **(a)** 206 IP genes with 410 patient survival associations in 33 cancer types, prioritised by their detection frequency in our elastic net framework (Y-axis). The IP genes associated with patient survival in glioblastoma (GBM) are labeled. **(b)** Catalogue of survival-associated IP genes in individual cancer types. Top: Bar plots of median univariate hazard ratios (HRs) and 95% confidence intervals. Bottom: Median expression values of IP genes. Z-transformed relative expression values of individual IP genes compared to all protein-coding genes are shown.

### High *GJB2* or *SCN9A* expression associates with poor survival of GBM patients

We focused on IP genes in GBM, a fatal form of brain cancer with an unmet need for target identification. Four high-confidence GBM IP genes were found, all of which were highly expressed in higher-risk patients: gap junction *GJB2* involved in non-syndromic hearing impairments (33), voltage-gated sodium channel *SCN9A* involved in pain sensation in the peripheral nervous system (34), aquaporin *AQP9* with roles in kidney cancer (35), and calcium-activated potassium channel *KCNN4* with roles in GBM (36,37). Given its the reported functions in GBM, *KCNN4* served as a positive control of our analysis. We confirmed the prognostic signals of these genes in multivariate analyses that accounted for patient age, sex, and the well-established GBM marker of *IDH1/2* mutation status (38) (**Figure 3a**). Besides GBM, *GJB2* and *SCN9A* expression profiles were associated with poor prognosis in low-grade glioma, kidney, and uterine cancer (**Figure S5b**). We selected *GJB2* and *SCN9A* for further studies in GBM.

**Figure 3.**
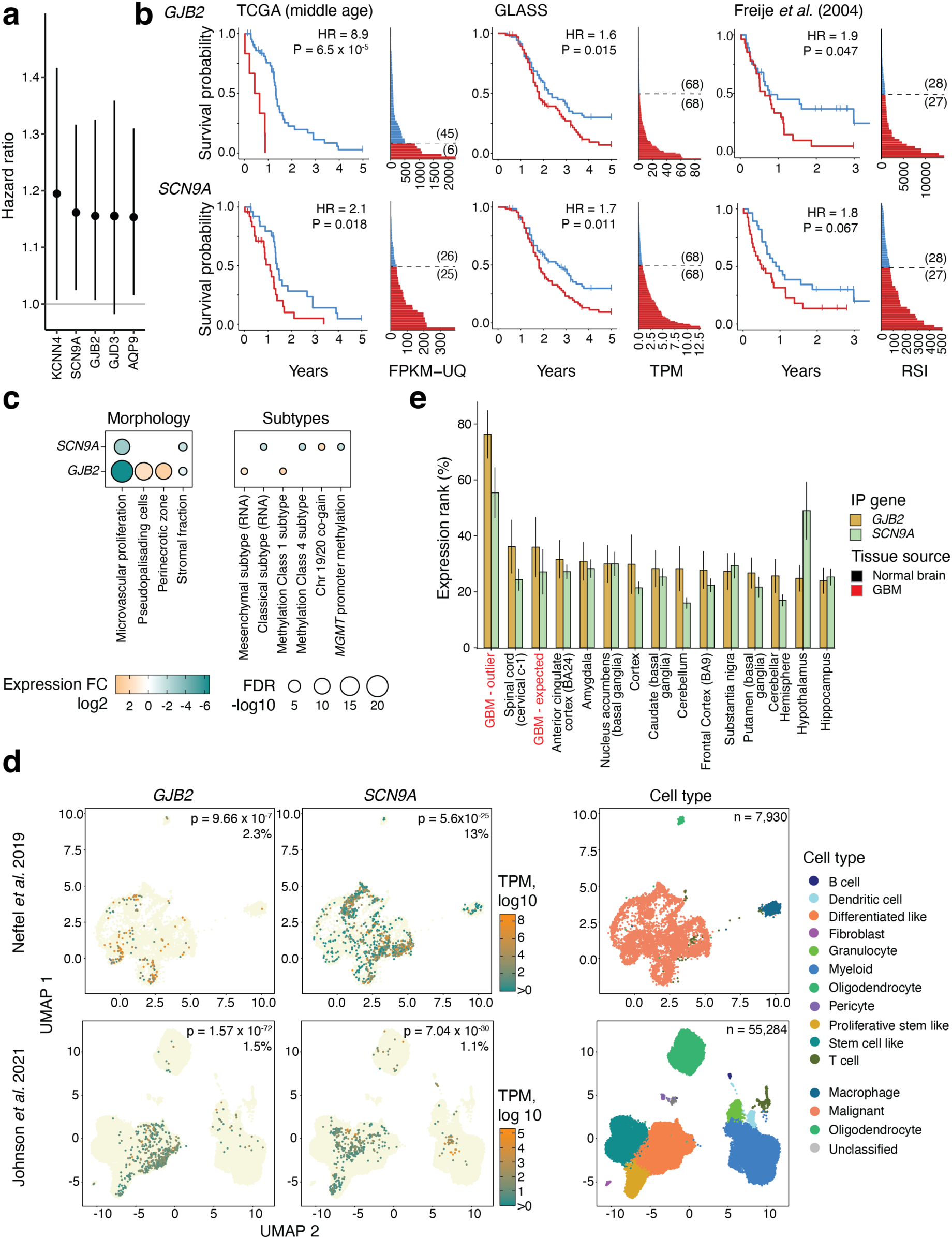
Survival associations and molecular features of *GJB2* and *SCN9A* in GBM. **(a)** Multivariate hazard ratios (HR) of IP genes prioritized in GBM. Median univariate HR is shown with 95% confidence intervals. **(b)** Kaplan-Meier plots of overall survival (OS) in GBM patients grouped by *GJB2* and *SCN9A* expression in TCGA (left) and two validation datasets (middle, right). Bar plots show gene expression in risk groups. Risk groups were determined by Tukey outlier analysis for *GJB2* and median dichotomisation for *SCN9A*. Wald *P*-values, Univariates HR values, and sample counts are shown. Patient age was included as a covariate in TCGA. **(c)** *GJB2* and *SCN9A* expression associates with GBM subtypes and anatomical regions. **(d)** *GJB2* and *SCN9A* expression in single glioma cells from two single-cell RNA-seq datasets shown as UMAP plots. Cells are colored by expression of *GJB2* (left) and *SCN9A* (middle). Cell type classifications of the original studies are shown on the right. **(e)** Comparison of *GJB2* and *SCN9A* expression in GBMs and normal brain samples from GTEx. Quantile-normalised median expression values with confidence intervals are shown (±1 s.d).

We examined the expression of *GJB2* and *SCN9A* in overall survival (OS) risk groups of GBM. For *GJB2*, GBM patients with high outlier expression had statistically worst prognosis (**Figure 3b, Figure S6a**). For *SCN9A*, the strongest association with poor prognosis was found by high gene expression by median-dichotomisation. Patient age was highlighted as a consistent prognostic factor in our ML-driven discovery of IP genes in GBM. To study the age component in detail, we analysed age-based tertiles of the GBM cohort and found strongest prognostic signals of *GJB2* and *SCN9A* in the middle age group (56 – 66 years, 51 patients), while the other tertiles showed attenuated signals (**Figure S6a**). The middle age group marks the greatest risk increase of presenting GBM, with twice the incidence rate compared to younger individuals and representing a third of the TCGA GBM cases within a decade of age (19).

We validated the survival associations of *GJB2* and *SCN9A* expression in two independent GBM cohorts, including 136 samples from the Glioma Longitudinal Analysis (GLASS) consortium (39) and 55 samples from a microarray-based dataset by Freije *et al*. (40) (**Figure 3b**). High *GJB2* or *SCN9A* expression based on median dichotomisation associated with poor prognosis in both datasets (*P* < 0.1; HR > 1.5), while weaker associations in samples with highly-elevated expression were also detected (**Figure S6b**). Associations with patient age and elevated IP expression should be confirmed in larger, better-powered cohorts. Collectively, the survival associations of *GJB2* and *SCN9A*, with elevated expression in high-risk GBMs, implicate these genes as potential targets to be investigated by functional experiments.

### *GJB2* and *SCN9A* expression is enriched in neoplastic cells and aggressive GBM subtypes

We sought to characterise *GJB2* and *SCN9A* in the contexts of GBM tumor regions and subtypes. First, we analysed anatomical datasets from the Ivy GBM Atlas, an anatomic transcriptional atlas of human GBM (41) (**Figure 3c**). Regions of microvascular proliferation showed reduced *GJB2* and *SCN9A* expression (log2FC < −2.9, FDR < 1.4 × 10^−3^). These represent a GBM hallmark comprising both resident endothelial cells and differentiated malignant cells (42). *GJB2* and *SCN9A* expression was lower in stromal fractions, which primarily include non-malignant fibroblasts (43). *GJB2* expression was higher near the necrotic centers of GBMs in pseudo-palisading and peri-necrotic zones. Thus, both *GJB2* and *SCN9A* are downregulated in anatomical regions characterized by less abundant tumor cells, while *GJB2* is upregulated in highly proliferative and motile regions of GBMs.

Next, we studied *GJB2* and *SCN9A* in the contexts of transcriptomic and methylation-based GBM subtypes and genomic alterations (21,44) (**Figure 3c**). *GJB2* expression was higher in mesenchymal GBM and related methylation subtype class-1 (log2 FC > 1.2, FDR < 0.05) (21). Patients with mesenchymal GBM have worse prognosis due to highly infiltrative and aggressive tumors (22). Lower *SCN9A* expression was found in classical GBM based on both classifications. Analysis of genomic alterations showed that *SCN9A* expression was associated with chromosome 19 and 20 co-gains which are found in many classical and some mesenchymal GBMs (21). Lower *SCN9A* expression associated with *MGMT* promoter methylation, an indicator of therapeutic response to temozolomide treatment (45).

We then determined the cell types expressing *GJB2* and *SCN9A* in GBMs using two single-cell transcriptomics datasets (46,47) (**Figure 3d**). Expression patterns of both genes were identified in subsets of neoplastic cells while their expression was undetectable in non-cancer cells, with a significant enrichment towards cancer cell fraction in both studies (*P* < 10^−6^, Fisher’s exact test). Among neoplastic cells, *GJB2* expression was higher in differentiated GBM cells and lower in proliferative stem-like cells. Higher *GJB2* expression in differentiated cells was characteristic of *IDH-*wild type GBMs with worse prognosis (46), while reduced *GJB2* expression and enrichment of stem-like cells was apparent in *IDH*-mutant gliomas with improved prognosis (38). Compared to *GJB2*, *SCN9A* expression was more uniform across neoplastic cell types. Besides neoplastic cells, *GJB2* expression was detected in myeloid cells while *SCN9A* was expressed in macrophages. Other non-cancer cells showed little or no expression of the two genes.

We compared *GJB2* and *SCN9A* expression in normal brain samples relative to their expression in GBMs using Tukey’s outlier analysis (**Figure 3e**). Expression profiles of 13 types of normal brain samples from 339 individuals were retrieved from the Genotype-Tissue Expression (GTEx) project (48). As expected, the GBMs classified as outliers showed significantly higher expression of *GJB2* and *SCN9A* than normal brain samples: *GJB2* ranked among the 24% most highly expressed genes in the outlier GBM group, while it ranked much lower (70%) in normal brain tissues and non-outlier GBMs (*P* = 1.5 × 10^−13^, Mann-Whitney U-test). Similarly, *SCN9A* expression was significantly higher in outlier GBMs compared to non-outlier GBMs and normal tissues (45% *vs*. 75%, *P* = 1.1 × 10^−9^). In contrast, *GJB2* and *SCN9A* expression in non-outlier GBMs was comparable to normal brain tissues.

Taken together, *GJB2* and *SCN9A* expression shows inter- and intratumoral heterogeneity in GBM. Their higher expression in malignant cell types and clinically relevant GBM subtypes implicate functional significance of these two IP genes in GBM.

### *GBJ2* or *SCN9A* knockdown deregulates proliferative and neural projection pathways

Next, we investigated the functional roles of *GJB2* and *SCN9A* using shRNA-mediated knockdown (KD) in a patient-derived GBM cell line (23,49). Transcriptome-wide profiling revealed dramatic changes induced by *GJB2* and *SCN9A* KD, with differential expression of 4,647 and 2,088 genes, respectively, including 640 common genes (absolute FC > 1.25, FDR < 0.05) (**Figure 4a**, **Figure S7a**). As expected, *GJB2* and *SCN9A* were significantly downregulated by KD (**Figure 4e**). To interpret these transcriptomic changes in the context of TCGA GBM tumors, we median-dichotomised patient samples by *GJB2* and *SCN9A* expression and uncovered hundreds of genes in differential expression analysis (**Figure S7b**).

**Figure 4.**
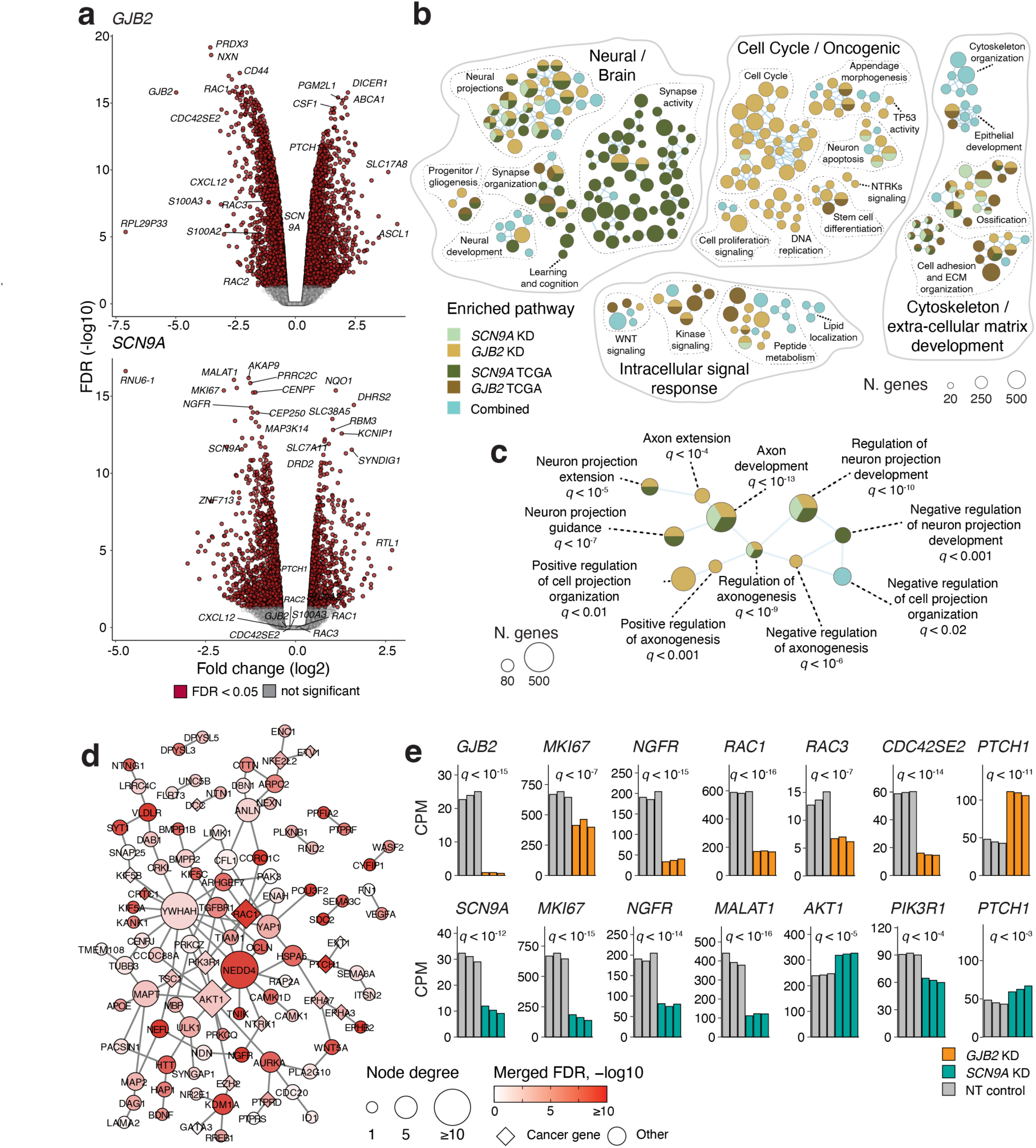
Transcriptomic profiling of GBM cells with *GJB2* and *SCN9A* knockdown (KD) indicates their roles in proliferative and neural projection pathways. **(a)** Genes differentially expressed genes in patient-derived GBM cells (G729) with *GJB2* and *SCN9A* KD (EdgeR, FC > 1.25, FDR < 0.05). **(b)** Pathway enrichment analysis of genes associated with *GJB2* and *SCN9A* expression (ActivePathways, FWER < 0.05). Differentially expressed genes from our KD experiments in GBM cells were jointly analysed with the genes identified in patient GBMs in TCGA. The enrichment map shows enriched pathways as nodes that are connected by edges into subnetworks if the pathways share many genes. Each pathway is colored by the transcriptomics dataset in which it was identified. **(c)** Subnetwork of neural projection pathways from panel (b). **(d)** Protein-protein interactions (PPIs) of genes from tunnelling nanotube pathways. Nodes show differentially expressed genes in GBM KDs of *GJB2* and *SCN9A* (panel (a)), that are connected by edges of high-confidence PPIs from the BioGRID database. **(e)** Differential expression of selected genes involved in tunneling nanotube pathways, mitosis, and signal transduction. Normalised gene expression values in counts per million (CPM) for *GJB2* or *SCN9A* KDs and non-targeting (NT) controls are shown. Q-values (FDR) from EdgeR are shown.

To define the genes and pathways associated with *GJB2* and *SCN9A* KD, we integrated the gene lists from patient-derived GBM cells and patient GBMs to identify jointly-enriched pathways using the ActivePathways method (50). We discovered four major functional themes with differential expression: cell proliferation, neural and brain development, signal transduction pathways, and cytoskeletal and extra-cellular matrix processes, with 350 significant processes and pathways in total (*FWER* < 0.05; ActivePathways) (**Figure 4b**, **Table S4**). These pathways included cancer hallmarks of cell proliferation, cell cycle deregulation, DNA replication, and neural apoptosis, as well as signal transduction cascades such as the WNT pathway. Cancer proliferation and invasion genes were downregulated, including proliferation marker gene *MKI67* with prognostic value in glioma (51), nerve growth factor receptor *NGFR* involved in GBM invasion (52), and long non-coding RNA *MALAT1* with tumor suppressive function in GBM (53) (**Figure 4e**). The enriched pathways were supported by multiple transcriptional signatures, indicating that target pathways of these two IP genes converge across our patient-derived GBM cell line models and patient tumors.

We focused on a group of neural projection processes that associated with both genes in our integrative pathway analysis. These included broader processes, such as regulation of neuron projection development (FDR = 1.4 × 10^−10^) and specific enrichments such as axonogenesis and dendrite development (**Figure 4c**). To prioritise individual genes in these pathways, we performed a network analysis that complemented the pathway analysis by examining interactions among genes. We reconstructed a protein-protein interaction (PPI) network that captured 102 of the differentially expressed neural projection genes, using interactomes from the BioGRID database (54) (**Figure 4d**, **Table S5**). The network highlighted *AKT1* and *PIK3R1* of the oncogenic PI3K/AKT signalling pathway involved in GBM (55), and the tumor suppressor patched homolog 1 (*PTCH1*) (56). *PTCH1* was upregulated in both *GJB2* and *SCN9A* KDs, while *AKT1* and *PIK3R1* were deregulated in *SCN9A* KD (**Figure 4d**). Overall, these results suggest that proliferation pathways are deregulated by our candidate genes.

Interestingly, we uncovered genes and pathways regulating tunnelling nanotubes (TNTs). TNTs, which are filipodia-like extensions between cells that enable cell-to-cell communication, promote tumor invasion, proliferation, and therapy resistance in GBM (57–60). Our pathway and network analyses highlighted two Rac family small GTPases (*RAC1, RAC3*) that were downregulated in *GJB2* KD cells, as well as the signalling adaptor *CDC42SE2* involved in TNT formation (61) (**Figure 4e**). Furthermore, the PI3K/AKT signalling pathway differentially expressed in *GJB2* KD GBM cells is implicated in TNT (57). Collectively, transcriptome-wide signatures of *GJB2* and *SCN9A* indicate their roles in proliferative and neural projection pathways in GBM. In particular, the TNT pathways deregulated in *GJB2* KD cells represent an intriguing avenue for further characterisation.

### GJB2 regulates the formation of tunneling nanotubes in GBM cells

To define the role of *GJB2* in TNTs, we investigated the cellular phenotypes of *GJB2* KD using two patient-derived GBM cell lines G797 and G729 (**Figure 5**). First, we determined the impact of *GJB2* KD on Rho GTPase pathway genes. *GJB2* KD significantly decreased the expression of RAC Rho GTPase genes *RAC1*, *RAC2*, and *RAC3*, in support of our findings from transcriptomics and pathway analyses (**Figure 5a**). Similarly, TNT-associated signaling adaptor gene *CDC42SE2* was downregulated in *GJB2* KD cells, while tumor suppressor *PTCH1* was significantly upregulated. Furthermore, *GJB2* KD reduced RAC1 protein expression (**Figure 5b**). Next, we monitored TNT dynamics using three patient-derived GBM cell lines (G411, G729, G797) (**Figure 5c**). While GBM cells formed robust TNT networks that connected different cells. we found a striking reduction in TNT lengths in all GBM cell lines upon *GJB2* KD (FC > 1.13, *P* < 0.05). Since TNTs can be formed from physical interaction of two filopodia in double filopodia bridges (62) and RAC1 is a critical regulator of filopodia formation (63,64), we investigated the role of *GJB2* on the dynamics of cell filopodia. Time-lapse imaging of membrane GFP-expressing G411 cells revealed that *GJB2* KD reduced the maximum extension length and lifetime of filopodia, while the extension rate and total number of filopodia remained unchanged (**Figure 5d**). Taken together, these results demonstrate that *GJB2* regulates filopodia dynamics and TNT formation in GBM cells.

**Figure 5.**
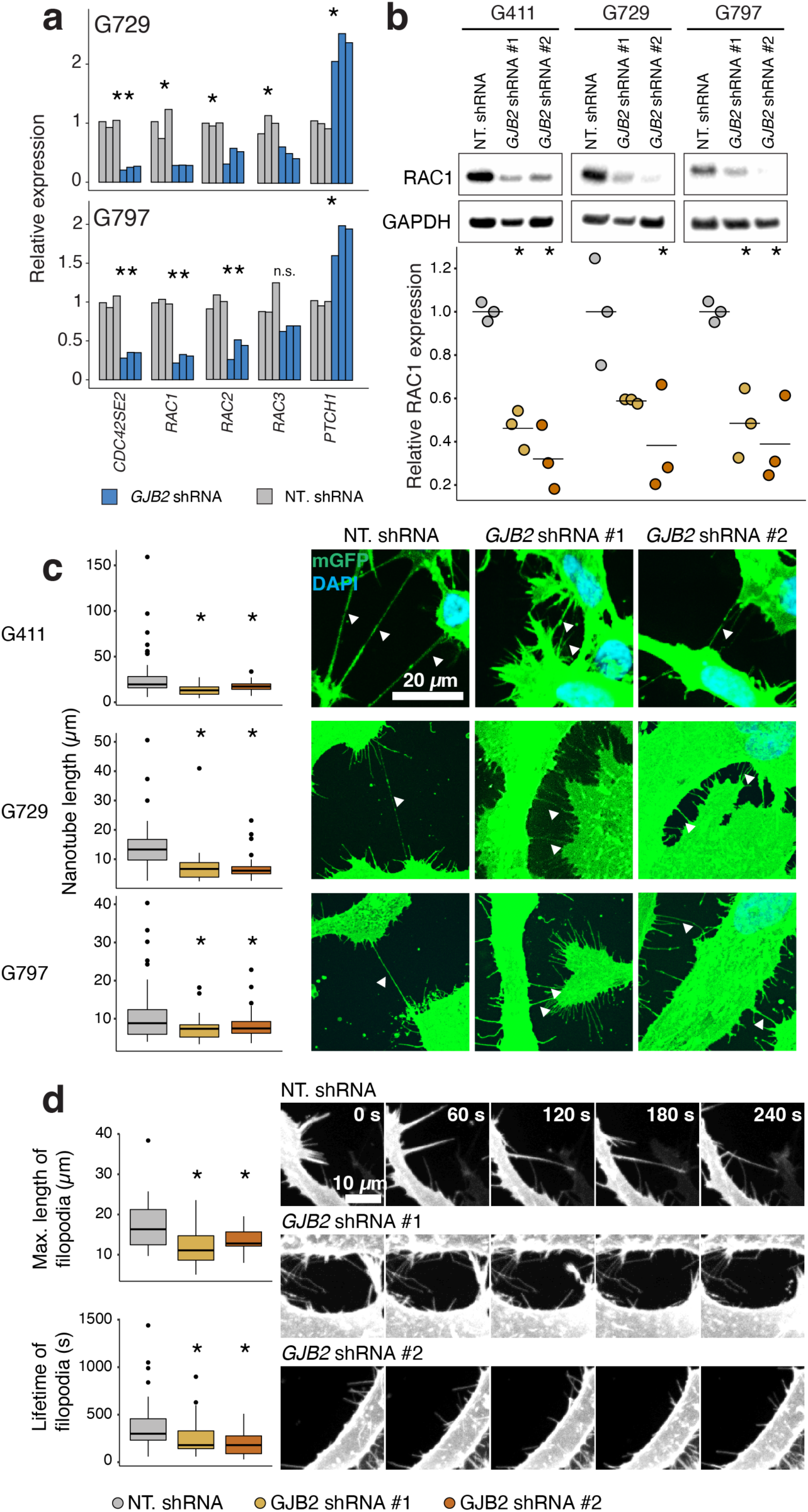
*GBJ2* knockdown in patient GBM cells perturbs tunnelling nanotube (TNT) formation. **(a)** TNT genes were dysregulated in *GJB2* KD cell lines. Comparison of mRNA expression of TNT pathway genes in *GJB2* KD GBM cells and non-targeted (NT) control cells measured using RT-qPCR. Relative gene expression values were normalised to control genes. P-values from FDR-adjusted t-tests are shown. **(b)** Protein quantitation of RAC1 in *GJB2* KD cells. Top: western blot of RAC1 and GAPDH (control) in three patient-derived cell lines targeted with two different *GJB2* shRNAs or NT controls. Bottom: relative expression of RAC1 in *GJB2* KD cells compared to NT controls from the western blot. Horizontal lines display the mean relative protein expression in each group, normalised in each cell line. **(c)** TNT projection lengths are reduced in *GJB2* KD cells. (Left) TNT lengths quantified from confocal microscopy images (right) of *GJB2* KD GBM cells tagged with membrane-targeted GFP and DAPI. **(d)** Timelapse images of membrane-GFP tagged GBM cells. Filopodia extension length and filipodia lifetime in *GJB2* KD GBM cells and NT control cells were measured for each cell at 30 second intervals over 180 intervals. All experimental results represent at least three independent replicates. P-values of t tests are shown (\**P* < 0.05, \*\**P* < 0.01).

### GJB2 and SCN9A promote GBM growth *in vitro* and *in vivo*

Finally, we investigated the role of GJB2 or SCN9A in regulating *in vitro* growth and *in vivo* tumorigenic potential of GBM cells. We studied three patient-derived mesenchymal GBM cell lines with high native expression of *GJB2* and *SCN9A* (23,24,49). First, we found that *GJB2* and *SCN9A* KD drastically reduced GBM cell viability (**Figure 6a**). Second, we evaluated the self-renewal capacity of GBM cells by determining their sphere forming ability using limited dilution assay (LDA). *GJB2* or *SCN9A* KD effectively abolished sphere formation (**Figure 6b**). Third, we investigated the roles of *GJB2* and *SCN9A* in regulating GBM growth *in vivo* (**Figure 6c-d**). We orthotopically injected luciferase-expressing G411 cells into immunodeficient NOD-SCID gamma mice. We monitored the survival of GBM-bearing mice and examined tumor growth using non-invasive bioluminescence imaging. *GJB2* and *SCN9A* KD markedly reduced tumor growth. Mice bearing *GJB2* or *SCN9A* KD GBM displayed significantly prolonged survival (*P* < 0.05, Wald test). Collectively, these results demonstrate that GJB2 and SCN9A promote GBM growth *in vitro* and *in vivo*, are consistent with the findings that GBM-relevant genes and pathways are altered by their deficiency (**Figure 4**), and establish GJB2 and SCN9A as functional regulators of GBM aggression.

**Figure 6.**
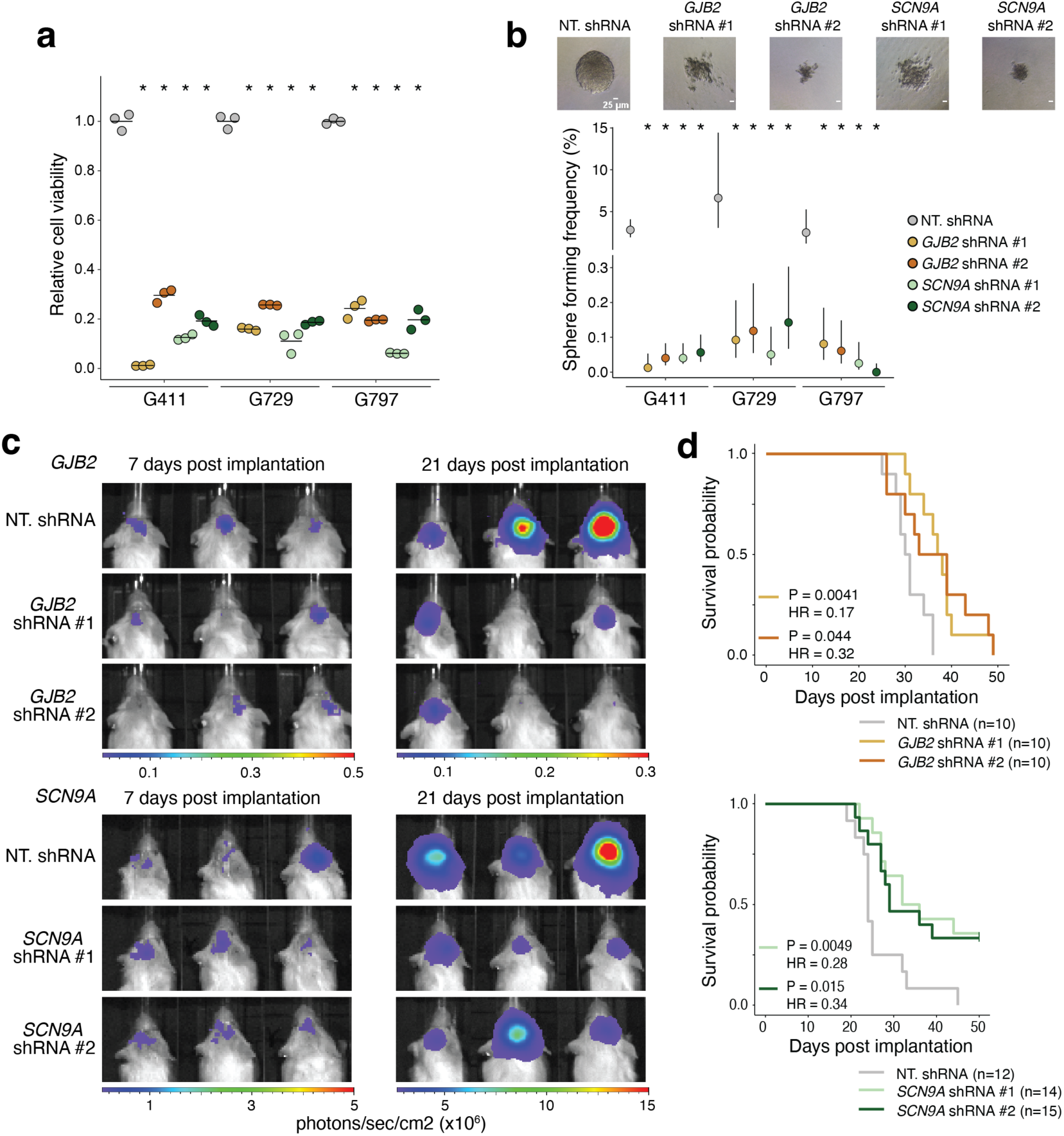
*GJB2* or *SCN9A* knockdown (KD) impairs GBM cell growth *in vitro* and *in vivo*. **(a)** *GJB2* or *SCN9A* KD reduces GBM cell viability *in vitro*. Cell viability was evaluated using an MTS assay in patient-derived GBM cell lines (G729, G797, G411). Horizontal bars show mean cell viability in each group, normalised to the mean of the controls (NT shRNA). FDR-adjusted P-values from t-tests are shown (all FDR < 0.01). **(b)** *GJB2* or *SCN9A* KD reduces sphere formation of GBM cells. Brightfield images and limited dilution assays (LDA) were performed on *GJB2*, *SCN9A* KD GBM cells and NT cells as controls. Sphere forming frequency was measured at the 14-day timepoint. Points show mean sphere formation frequency for each group of six replicates and the vertical lines show the full range of measurements. FDR-adjusted P-values from Mann-Whitney U-tests are shown (all FDR < 0.001). **(c)** *GJB2* or *SCN9A* KD impairs tumor growth in patient-derived GBM xenografts in mice. Bioluminescence imaging was performed on mice following patient-derived implantation of KD and NT cells of GBM cells (G411). Radiance was measured 10 minutes after injection with 100 mg/kg luciferin on the IVIS Spectrum system. **(d)** *GJB2* or *SCN9A* KD improves mouse survival in patient-derived GBM xenografts. Mice with KD and NT GBM cell xenografts (G411) were monitored for survival for 50 days and visualised as Kaplan-Meier plots. Survival analysis was performed independently for each shRNA. Wald *P*-values and Cox proportional-hazard (CoxPH) hazard ratios (HR) are shown.

## DISCUSSION

Ion channels comprise a large class of drug targets. More than 15% of U.S. FDA-approved drugs target ion channels to treat a variety of human diseases, such as diabetes, hypertension, and neurological disorders (5–7). However, the expression patterns or functional roles of ion-permeating proteins are understudied in cancer. In this study, we delineated the transcriptomic landscape of IP genes in human cancer. We discovered that IP genes (including ion channels, ion transporters, and gap junctions) are highly upregulated in subsets of cancer samples at a frequency and magnitude that significantly exceed most protein-coding genes. This phenomenon is shared among dozens of major cancer types, thereby revealing a fundamental characteristic of cancer. On average, each cancer sample displays dozens of IP genes upregulated at levels significantly exceeding their physiological range. Integrative analysis of tissue-specific expression patterns and associations with patient survival outcomes provides a comprehensive catalogue of candidate genes with potential roles in mediating bioelectrical signalling in cancer. This catalogue serves as a useful resource for interrogating candidates for identifying biomarkers, validating therapeutic targets, and repurposing approved drugs that act on IP proteins.

*GJB2* and *SCN9A* are implicated in monogenic diseases with emerging implication in cancer. *SCN9A* functions in signal transduction in neurons, such as nociceptor pain signaling (65). *GJB2* encodes a gap-junction protein (connexin) whose autosomal recessive allele causes deafness in Asian populations (66). High *GJB2* expression is associated with worse prognosis in GBM and LGG, suggesting common mechanisms in high- and low-grade gliomas. *GJB2* and *SCN9A* have been linked to invasion and proliferation in prostate (67), lung (68), gastric (69), and breast cancer (70), as well as metastasis (71,72) and worse prognosis in several cancer types (70,73–75). In GBM, however, the phenotypic and prognostic aspects of *GJB2* and *SCN9A* have not been characterised to date. We selected *GJB2* and *SCN9A* as high-priority target genes in GBM due to their significant associations with patient survival identified in our machine learning analyses. *GJB2* or *SCN9A* KD led to profound transcriptional dysregulation that disrupted proliferative and neural projection pathways in patient-derived GBM cell lines. Notably, *GJB2* KD affected TNT pathways that control intercellular communications within GBM. The lengths of TNTs and filipodia were disrupted by *GJB2* depletion in GBM cells, possibly via reduced RAC1 expression. We demonstrated that reduced expression of either *GJB2* or *SCN9A* strongly impaired GBM cell viability *in vitro* and *in vivo*. Further, both genes show intratumoral variation in GBM and are predominantly expressed in malignant cell types. Collectively, these data establish our top-listed genes as functional regulators of GBM aggression.

GBM networks are comprised of multicellular connections between tumor cells and neurons, astrocytes, and other cells of the tumor microenvironment that are indispensable for proliferation, invasion, metabolic rewiring, and therapy resistance (76–80). Different connections have been characterised: tumor microtubes (TMTs) are over 500 µm in length, last for days, and consist of gap junction connections, whereas TNTs are shorter than 100 µm, last for hours, and are mostly open-ended with few connections (80). *GJB2* KD GBM cells showed reduced expression of RAC small GTPases (RAC1-3) and the CDC42 effector *CDC42SE2* that are involved in TNTs (81,82), while no expression changes were found for previously identified TMT-regulating genes (76,83), suggesting that *GJB2* function may be specific to TNTs. Gap junction proteins, such as connexin 43 (*GJA1*), mediate intercellular electrical signaling through TNTs (84,85). Electrical coupling of GBM cells through gap junctions is required for tumor growth (77). Determining the subcellular localisation of GJB2 and its role in electrical conductance is needed to establish GJB2 as a direct regulator of TNTs. We found that RAC Rho GTPases and actin regulators were downregulated in GBM cells upon *GJB2* KD. RAC1 is a major regulator of actin dynamics (62,63), and TNTs form from actin-rich membrane protrusions (58,61,79). Thus, our results suggest that GJB2 may also affect TNTs indirectly through RAC1 and other actin regulators. This is consistent with previous observations in HeLa cells, where *GJB2* overexpression increased RAC1 activation (86). Further studies are required to determine how GJB2 regulates RAC1.

In sum, our study not only reveals that prominent activation of IP genes is associated with tumor heterogeneity and patient outcomes but also establishes specific IP genes with oncogenic roles in GBM. The global IP gene alterations indicate that ionic flux-mediated bioelectrical signalling via aberrant ion permeome activity is a potential pan-cancer hallmark.

## Methods

### TCGA transcriptomics data and patient clinical information

Bulk tumor RNA-seq data and patient clinical information of the TCGA PanCanAtlas project (25) were collected from the Genomic Data Commons. RNA-seq data in FPKM-UQ (fragments per kilobase million, upper quartile) were used unless specified otherwise. In cases with multiple tumor samples per patient, we selected the sample with the first barcode. Control samples and tumor samples lacking survival or RNA-seq data were removed. We analysed cancer types with at least 50 samples, with 9352 samples of 29 cancer types in total, including 150 GBMs. Breast cancer (BRCA) subtypes were analysed separately (luminal A, luminal B, *HER2* positive, basal-like, normal-like), using annotations from the R package *TCGABiolinks* (44), resulting in distinct 33 cancer types. In gliomas (GBM, LGG), *IDH1/2* mutation status from *TCGABiolinks* was included as a clinical covariate. All GBM samples from TCGA we analysed were *IDH1/2* wildtype or unclassified. Data analysis was performed in Python (3.9.11) using custom scripts. Unless stated otherwise, statistical tests were performed using the *stats* package from the *Scipy* software library. The *ggplot2* R package was used for visualisations (R 4.1.3, *ggplot* 3.4.0).

### Ion permeome genes

Drug targetable ion permeome (IP) genes were retrieved from the Guide to Pharmacology (GtP) database (downloaded June 6, 2022) (26). IP genes included the classifications of voltage-gated ion channels (ICs), ligand-gated ICs, and other ICs. As controls, we studied two drug target families: kinases and G-protein coupled receptors (GPCRs). GPCRs were obtained from GtP. Kinases were retrieved from the UniProt database (pKinFam.txt, downloaded Sept. 15, 2022) (87) and intersected with the list of enzymes in GtP. Genes lacking RNA-seq data in TCGA were excluded. In total, 276 IP genes, 391 GPCRs, and 505 kinases were included.

### Clustering cancer samples by IP gene expression

An unsupervised analysis of cancer samples using IP gene expression as features was performed using standardised, log1p-transformed FPKM-UQ expression values. The Uniform Manifold Approximation and Projection (UMAP) python package (88) with default parameters was used for dimensionality reduction. Cancer samples were visualised in the first two UMAP dimensions and colored by cancer type.

### Highly elevated expression of IP genes

We identified IP genes with highly elevated expression using Tukey’s outlier analysis (27). Each cancer type and IP gene was analysed separately. A cancer sample was considered to have highly-elevated (outlier) expression of a given IP gene if its expression exceeded the 75^th^ percentile of its expression across all samples of the given cancer type by 1.5-fold the interquartile range (25-75%). Otherwise, the sample was classified as having an expected expression range. We computed expression fold-change (FC) values, comparing the cancer samples having highly elevated and expected expression of IP genes, as the ratio of median expression values of the two groups. Switch-like IP genes were annotated separately. Switch-like IP genes had zero expression in most samples (*i.e.,* median zero) and fewer cancer samples with non-zero expression.

### Statistical analysis of elevated expression of IP genes

To evaluate the significance of elevated IP expression in cancer, we performed control analyses using (i) all protein-coding genes, and major classes of drug targets including (ii) GPCR genes, and (iii) kinase genes, as. The control gene sets were downsampled with replacement to match the count of IP genes (276) over 10,000 iterations. Each cancer type was analysed separately. Fractions of outlier cancer samples and median FC values from these iterations were used as controls to evaluate the cohort frequency and magnitude (fold-change) of IP gene upregulation in cancer. Cohort frequency and FC were visualised as 2D density plots for individual cancer types and the pan-cancer cohort. For control gene sets, representative iterations corresponding to median fold-change values were shown.

### Identifying survival-associated IP genes

To find IP genes significantly associated with patient survival, we used a machine learning framework based on Cox proportional hazards (CoxPH) elastic net models and bootstrap analysis adapted from our previous work (28). Log1p-transformed expression profiles of 276 IP genes were used as model features. Cancer types were analysed separately, with a model response of either overall survival (OS) or progression-free survival (PFS), as recommended previously (89) (**Table S1**). In each cancer type, IP genes with detectable expression were included (mean FPKM-UQ > 1 across all samples). IP genes were prioritised over 1,000 iterations of elastic net survival regression models fit on random subsets of 80% of samples. At each iteration, feature pre-selection selected a subset of IP genes that associated with survival in univariate CoxPH regression (Wald test; *P* < 0.1). These were fitted using the Python package *CoxPHFitter* from the *lifelines* library. A multivariate CoxPH model was then fitted with pre-selected genes as features and patient survival as response. Clinical variables were also included as features to evaluate the complementarity of IP genes (patient age and sex, tumor grade and stage; and *IDH1/2* mutations for GBM and LGG (38)). We selected the best-performing penalty (α) using a grid-search with 5-fold cross validations using the *GridSearchCV* package from *sklearn*. Following elastic net regularisation, model features (*i.e.*, IP genes and clinical variables) with non-zero coefficients were recorded. After all iterations, we selected the final IP candidate genes and clinical variables that were identified as features in the regularised CoxPH models (>50% iterations). To derive hazard ratios (HR) and 95% confidence intervals for the selected IPs, univariate CoxPH models using all samples were used. Elastic net training, regularisation and parameter evaluation was conducted using the *CoxnetSurvivalAnalysis* package from the *sksurv* library, with a fixed L1-ratio hyperparameter (λ = 0.5). Finally, to confirm that our approach was calibrated, we repeated the IP prioritisation workflow using 1,000 simulated datasets generated by randomly shuffling patient survival outcomes while maintaining true IP gene expression profiles. We compared the results of these simulated datasets to the true datasets. As expected, simulated data revealed significantly fewer and lower-confidence IP genes compared to true datasets (**Figure S3a**).

### Additional survival analyses of IP genes in GBM

In GBM, we focused on the five prioritised IP genes. Further vetting included extended multivariate CoxPH models with patient age, sex, and *IDH1/2* mutations as features. Based on HRs and Wald *P*-values, we selected four IP genes (*GJB2*, *SCN9A, KCNN4*, *AQP9*) and excluded *GJD3* due to sub-significant survival association and HR in multivariate models. To evaluate survival associations, GBM samples were split into two groups using median-dichotomisation and outlier-based (Tukey) dichotomisation of IP expression. Survival associations were evaluated using CoxPH regression separately for the discovery data (TCGA) and two external validation datasets (see below). Survival associations of *GJB2* and *SCN9A* were the strongest in the middle age group of TCGA GBMs (55 – 66 years), potentially explained by the age variable that was the strongest feature identified in our analysis.

### External validation of survival associations

We used additional GBM transcriptomics datasets to validate the survival associations of *GJB2* and *SCN9A*. First, we studied 136 primary GBMs profiled in the Glioma Longitudinal Analysis (GLASS) Consortium (39), after excluding recurrent GBMs, duplicate samples per patient, and samples used in TCGA. All GLASS samples were *IDH1/2* wildtype. GLASS RNA-seq data were available as transcripts per million (TPM) units. Second, we used the microarray dataset by Freije *et al.* (40) (GEO accession: GSE4412) with 55 grade-IV gliomas for which *IDH1/2* mutation status was unavailable. Relative fluorescent units (RFI) of gene expression were exponentially transformed to approximate normal distributions. In case of multiple cancer samples per patient, the alphabetically first sample was selected. We also performed survival analyses with covariates as described above. No significant associations with patient age were found, potentially due to smaller sample sizes or cohort composition.

### Candidate gene expression correlations with clinical, immune, and micro-environment features

To explore the potential roles *of GJB2 and SCN9A* in GBM, we asked how their expression associated with molecular and pathological features of GBMs, including longitudinal expression profiles from the *GLASS* project (39) and anatomical expression data of the Ivy Glioblastoma Atlas (41). The GBM subtypes based on DNA methylation and gene expression patterns were acquired from TCGA (21). Immune cell infiltration and immune-based cancer subtypes of TCGA samples were obtained from Thorsson *et al.* (90). Molecular features of TCGA tumors, including recurrent somatic mutations, copy number alterations, and clinical subtypes were obtained from *TCGABiolinks* (44). To associate features with the candidate genes, gene expression in samples with and without features were compared using two-tailed Mann-Whitney U-tests. For tumor subtypes and other multi-class features, we compared samples of a given subtype with samples of all other subtypes combined. Immune cell infiltration (ICI) profiles from CIBERSORT were first median dichotomised into two equal subsets of GBMs (high *vs.* low ICI). IP gene expression was compared between the resulting groups. Multiple testing correction was applied within each analysis and significant findings were reported (FDR < 0.05).

### IP gene expression in tumors and normal brain tissues

To compare *GJB2* and *SCN9A* expression in tumors and normal tissues, we analysed GBMs from TCGA and normal brain tissues from the Genotype-Tissue Expression (GTEx) dataset (version phs000424.v9, downloaded Oct. 5, 2022) (48). Expression data (TPM) for 3326 samples from 399 patients were obtained from the GTEx data portal. For improved comparison, RNA-seq data in TCGA and GTEx were then rank-normalised across all protein-coding genes. Expression ranks of *GJB2* and *SCN9A* were compared between 13 types of GTEx brain tissues and two subsets of GBMs (*i.e.*, GBMs with highly-elevated outlier IP gene expression, and GBMs with expected expression). The mean expression ranks were shown with +/- one standard deviation (s.d.) for each tissue type. Ranks of tissue types were compared using two-tailed Mann-Whitney U-tests and multiple testing correction (FDR) was applied.

### Expression of candidate IPs in GBM at the single-cell level

We studied *GJB2* and *SCN9A* expression in single-cell RNA-seq data of GBM samples from two studies: Neftel *et al.* (47) (7,930 cells, *IDH1/2* wildtype GBMs) and Johnson *et al.* (46) (55,284 cells, *IDH1/2* wildtype and mutant GBMs). The UMAP method was applied to log1p-transformed TPM values. Cells were coloured by expression of *GJB2* or *SCN9A* (log10 TPM). Cell type annotations were retrieved from the original studies.

### Patient-derived GBM samples and cell culture

GBM cells for functional experiments were obtained following informed consent from patients. Experiments were in accordance with the Research Ethics Board at The Hospital for Sick Children (Toronto, Canada). Access to pathological data was obtained from the institutional review boards. GBM stem cell lines (G797, G729, G411), which were established from mesenchymal GBMs, were cultured using previously established protocols (91) including serum-free NS cell self-renewal media (NS media) consisting of Neurocult NS-A Basal media, supplemented with 2 mmol/L L-glutamine, hormone mix (in house equivalent to N2), B27 supplements, 75 mg/mL BSA, 2 mg/mL Heparin, 10 ng/mL basic FGF and 10 ng/mL human EGF. GBM cell lines were grown adherently on culture plates coated with poly-L-ornithine and laminin and maintained in 37°C tissue culture incubator with 5% CO2. All cell lines were regularly checked for mycoplasma infections by DAPI staining.

### Knockdown of GJB2 and SCN9A in GBM cells

Knockdown experiments in GBM cell lines (G411, G729, G797) were performed using lentivirus-mediated shRNAs. KDs were repeated for three replicates per cell lines. KD efficacy of *GJB2* and *SCN9A* was validated using RT-qPCR. Human pLKO.1 lentiviral shRNA target against *GJB2* or *SCN9A* and pLKO.1-TRC-control vector were obtained from Dharmacon. Viral transduction was performed in antibiotics-free culture medium for 24 hours. The following shRNA mature antisense sequences were used: *GJB2* #1: GTCTTCACAGTGTTCATGATT; #2: GAACGTGTGCTACGATCACTA. *SCN9A* #1: GCCCTCATTGAACAACGCATT; #2: GCTGATTTGATTGAAACGTAT.

### RNA-seq profiling of GJB2 and SCN9A knockdowns

RNA-seq data was generated in the patient-derived GBM cell line G729 with shRNA targeting *GJB2*, *SCN9A*, or non-targeting (NT) shRNA controls, as described above. Total RNA was collected 4 days post lentiviral shRNA transduction using RNeasy Plus Mini Kit (Qiagen #74134). Lentiviral transduction and RNA extraction were performed in triplicates. RNA integrity number (RIN) was determined using Agilent Bioanalyzer. All samples had RIN > 9.8. Library preparation was performed using NEBNext Ultra II Directional polyA mRNA Library Prep Kit. Sequencing was performed on Illumina NovaSeq 6000 with 30 million paired end reads per sample at 100 bp read length.

### Processing and data analysis of RNA-seq data of KD cell lines

We aligned RNA-seq reads to the reference human genome HG19 (GRCh37.p13) from the GENCODE, for better consistency with the TCGA dataset. Reads were mapped to the transcriptome using Rsubread (92) with default settings. Differential gene expression analysis of *GJB2* and *SCN9A* KD cells was conducted on raw read counts. Three replicates treated with IC-targeting shRNAs for each IP gene were compared to three control replicates treated with NT shRNAs, using the *EdgeR package* in R (93). First, lowly expressed genes were removed (mean count < 1). Second, we selected significantly differentially expressed genes with an absolute fold change of at least 1.25 (abs log2 fold change *flc* > 0.32) using the *glmTreat* method of EdgeR. P-values from *EdgeR* were corrected for multiple testing and significant genes were selected (FDR < 0.05). Individual gene expression values were visualised using counts per million (CPM).

### Transcriptomics analysis of TCGA data for GJB2 and SCN9A

To integrate the transcriptomics data from our KD experiments with patient tumor data, we detected the genes associating with *GJB2* and *SCN9A* expression in the GBMs in TCGA. Raw RNA-seq counts were obtained from *TCGABiolinks* (44) and lowly expressed genes were filtered (mean count < 1). TCGA GBMs were split into two groups based on the median gene expression separately for *GJB2* or *SCN9A*. Differential gene expression analysis was used to compare the groups using *glmTreat* from the *EdgeR* package (93) and significant genes were selected (abs FC ≥ 1.25, FDR < 0.05).

### Integrative pathway enrichment analysis

We performed an integrative pathway enrichment analysis to identify pathways and processes jointly associated with *GJB2* and *SCN9A* expression in our cell line KD experiments and GBMs in TCGA. We used the *ActivePathways* method (94) with a matrix of P-values representing differential expression estimates of all protein-coding genes in the four contrasts (*GJB2* and *SCN9A*; both in cell lines and TCGA). Gene sets of biological processes from Gene Ontology and molecular pathways from Reactome were downloaded from the gProfiler web server (Jan. 13 2023) (95). Gene sets with 50 – 500 genes were used. All protein-coding genes measured in RNA-seq datasets were used as the background set. Genes with low expression (mean count < 1) were deprioritised prior to the analysis by setting their P-values to 1.0. The resulting pathways were corrected for multiple-testing and significant results were selected using default settings (Holm family-wise error rate (*FWER)* < 0.05). The enrichment map of pathways and processes was created using the EnrichmentMap app in Cytoscape standard protocols (96). The major functional themes were organised manually.

### Protein-protein interactions of neuron projection pathways

We constructed a PPI network of the differentially expressed genes in *GJB2* and *SCN9A* KD experiments that were annotated in neural projection pathways. First, we selected a subset of GO processes related to neuron projection from our pathway enrichment analysis (**Table S5**). Of those pathways, we selected the genes that were differentially expressed in at least one KD experiment. All human PPIs were downloaded from the BioGRID database (54) (version 4.4.217, July 22, 2022) and filtered to include only high-confidence PPIs found in at least two studies (PubMed IDs). Self-interaction PPIs were excluded. The PPI network was then limited to the neural projection genes defined above and visualised using Cytoscape (96). Proteins in the network were prioritised by node degree (node size) and differential expression in KD experiments (node color; merged FDR values using Brown’s method). We highlighted known cancer genes of the COSMIC Cancer Census database (97).

### Mouse xenograft experiments

We performed orthotopic mice xenografts of patient-derived GBM cell lines using female NOD SCID gamma /J#5557 immunodeficient mice aged eight weeks. Mice were housed under aseptic conditions with filtered air and sterilised food, water, bedding, and cages. Animal procedures followed the Animals for Research Act of Ontario and the Guidelines of the Canadian Council on Animal Care, as approved by the Centre for Phenogenomics (TCP) Animal Care Committee (protocol 23-0288H). GBM cells (G411) were transduced with lentiviral vector pBMN (CMV-copGFP-Luc2-Puro, Addgene plasmid #80389, a gift from Magnus Essand) expressing GFP and firefly luciferase under the control of cytomegalovirus (CMV) promoter. GFP+ cells were sorted by fluorescence-activated cell sorting (FACS). G411 GFP-Luc2 cells were transduced with NT, *GJB2* or *SCN9A* lentiviral shRNAs for 24 hours. Two days post transduction, cells were injected into mice. Mice were anesthetised using gaseous isoflurane and immobilised in a stereotaxic head frame. The skull of the mouse was exposed, and a small opening was made using sterile dental drill (Precision Guide) at 1 mm lateral and 2 mm posterior to bregma. At this location, 2000 cells were injected with a Hamilton syringe 2.5 mm deep at a rate of 1 µL/min using a programmable syringe pump (Harvard Apparatus). *GJB2* KD and *SCN9A* KD xenografts were performed in separate batches. For *GJB2* KD, 10 mice were used for each of NT, shRNA #1, and shRNA #2 groups. For *SCN9A* KD, 12 mice were used for NT, 14 mice for shRNA #1, and 15 mice for shRNA #2. All procedures were carried out under sterile conditions. Kaplan-Meier curves were generated to compare the survival of mice in the groups.

### In vivo bioluminescence imaging

*In vivo* bioluminescence imaging was performed using the Xenogen IVIS Lumina System coupled with LivingImage software for data acquisition. Mice were anesthetised using gaseous isoflurane and imaged 10 minutes after intraperitoneal injection of 100 mg/kg luciferin.

### Cell viability, limited dilution assay, tunneling nanotube and filopodia imaging

Cell viability was determined using CellTiter 96 AQueous One Solution Cell Proliferation Assay (Promega), utilising MTS reagent, which produces a colored formazan dye when metabolised by NAD(P)H-dependent dehydrogenase enzymes (98). GBM cells were plated at 1000 cells per well in poly-L-ornithine and laminin coated 96 well plates and transduced with a 50-fold dilution series of lentiviral shRNA for 24 hours. 7 days post transduction, CellTiter 96 AQueous One Solution Cell Proliferation Assay was performed according to manufacturer protocol. Formazan dye absorbance was read using a microplate reader (Molecular Devices). Differences in cell viability were tested using a two-tailed independent-sample t-tests. Limited dilution assay was performed by plating GBM cells in a serial dilution ranging from 2000 to 3 cells per well on round-bottom 96-well plates in six biological replicates. The numbers of wells with spheres were quantified seven days after plating and data were analysed by Extreme Limiting Dilution Analysis (ELDA) software (59), which calculates the frequency of sphere forming cells and the differences between groups using Chi-square tests. GBM cells were transduced with lentiviral membrane-GFP (exact construct, Addgene #22479). GFP+ cells were sorted by FACS. Membrane-GFP expressing GBM cells were transduced with lentiviral shRNA (MOI > 1). For imaging of tunneling nanotubes at 4 days post transduction, cells were fixed with 4% paraformaldehyde at room temperature for 20 minutes, then stained with DAPI. Images were acquired on Quorum Spinning Disk confocal microscope with 63x/1.4NA objective. For live imaging at 4 days post transduction, cells were imaged every 30 seconds for 90 minutes. Tunneling nanotube and filopodia length was quantified using the ImageJ software. The File Name Encryptor plugin was used to blind image file names, and the Line Measure tool was used to measure nanotube lengths. Significant differences in nanotube measurements were tested for using two-tailed Mann-Whitney U-tests.

### RNA extraction, reverse transcription, and RT-qPCR

Total RNA was collected 4 days post lentiviral shRNA transduction using GENEzol TriRNA Pure Kit (Geneaid #GZX200). RNA concentration was measured using NanoDrop 1000 Spectrophotometer, and 1 µg of RNA was reverse transcribed to cDNA using SensiFAST cDNA Synthesis Kit (Bioline #65054). qPCR reactions were set up using PowerUp SYBR Green Master Mix (Applied Biosystems #A25742) and real-time detection and quantification of cDNAs was performed on the Viia7 Cycler (Applied Biosystems) with 40 cycles of amplification. Viia7 System Software (Applied Biosystems) was used to determine Ct values with automatically set thresholds. Gene expression was normalised to *GAPDH* and analysed using the ΔΔCt method. The following RT-qPCR primers were used (h, human): hGAPDH, 5’-CTCCTGCACCACCAACTGCT-3’ (forward), 5′-GGGCCATCCACAGTCTTCTG-3′ (reverse); hRAC1, 5’-CGGTGAATCTGGGCTTATGGGA-3’ (forward), 5’-GGAGGTTATATCCTTACCGTACG-3’ (reverse); hRAC2, 5’-CAGCCAATGTGATGGTGGACAG-3’ (forward), 5’-GGAGAAGCAGATGAGGAAGACG-3’ (reverse); hRAC3, 5’-ACAAGGACACCATTGAGCGGCT-3’ (forward), 5’-CCTCGTCAAACACTGTCTTCAGG-3’ (reverse); hCDC42SE2, 5’-GGATCAGGAGACCTGTTCAGTG-3’ (forward), 5’-CCTTCGTATCCACGAGCTGCAT-3’ (reverse); hPTCH1, 5’-GCTGCACTACTTCAGAGACTGG-3’ (forward), 5’-CACCAGGAGTTTGTAGGCAAGG-3’ (reverse).

### Protein extraction and western blots

We quantified the protein levels of RAC1 in *GJB2* KD GBM cells. Total protein was extracted from multiple cell cultures (G411, G729, G797) using the RAC1 Activation Assay Biochem Kit per manufacturer’s instructions (Cytoskeleton Inc, #BK035-S) five days post transduction with lentiviral non-targeting or *GJB2* shRNA. All protein lysates were homogenised for 20 minutes at 4 degrees Celsius, then centrifuged at 4 degrees Celsius and 14,000 rpm for 10 minutes. 10 mg of protein samples were resolved on a 10% Bis-Tris gel (Invitrogen, #NW00102BOX) at 200 V in MES running buffer (Invitrogen, #B0002). The proteins were transferred onto a PVDF membrane (Millipore, #IPVH0001) and blocked with 5% BSA in 0.1% Tween-20 in TBS. Membranes were incubated overnight at 4 degrees Celsius in primary antibodies diluted in the blocking solution. Immunoreactive bands were visualised using HRP-conjugated secondary antibodies (Cell Signaling Technology), followed by chemiluminescence with ECL-plus Western Blotting Detection System (Amersham, #RPN2232). Chemiluminescence was imaged and analysed using Molecular Imager VersaDoc MP4000 system (Bio-Rad). The primary antibodies used were mouse anti-Rac1 (1:500, Cytoskeleton Inc, #ARC03) and rabbit anti-GAPDH (1:3000, Cell Signaling #2118S). Experiments were performed in three biological replicates. Significant differences in protein expression were analysed using two-tailed independent-sample t-tests.

### Statistical analyses of patient-derived cell lines and mice

No statistical methods were used to predetermine sample sizes in validation experiments. Statistical analyses were completed after the experiments without interim data analysis. No data points were excluded. All data were collected and processed randomly. Each experiment was successfully reproduced at least three times and the experiments were performed on different days.

## Acknowledgements

This work was supported by the New Investigator Award of the Terry Fox Research Institute (TFRI) to J.R., the Canadian Cancer Society (CCS) Innovation Grant to J.R. and X.H., the Canadian Institutes of Health Research (CIHR) Project Grant to J.R., and the Investigator Award to J.R. from the Ontario Institute for Cancer Research (OICR). Funding to OICR is provided by the Government of Ontario. A.T.B. was supported by the Ontario Graduate Scholarship (OGS). H.-K.M. was supported by OGS and the SickKids RestraComp Scholarship. The results shown here are in whole or part based upon data generated by the TCGA Research Network: https://www.cancer.gov/tcga.

## Author contributions

A.B. performed the data analysis. H.M. performed the experiments. A.B., H.M., X.H., and J.R. interpreted the data and wrote the manuscript. J.R. supervised the data analysis. X.H. supervised the experiments. K.G., A.F., I.D., and M.B. contributed to data analysis. W.D., H.Z., J.C. and X.C. contributed to experiments. P.B.D. acquired clinical GBM samples and cell lines. J.R. and X.H. conceived and supervised the project. All authors reviewed the manuscript and approved the final version.

## Supplementary tables

**Table S1.** Cancer types and number of samples analyzed.

**Table S2.** Druggable IP genes used in this study.

**Table S3.** Prioritized IP genes associated with survival outcomes.

**Table S4.** Complete list of pathways enriched from cell lines and TCGA differential genes expression.

**Table S5.** Neural projection pathways enriched from cell lines and TCGA differential genes expression.

## Notes

### Competing Interest Statement

The authors have declared no competing interest.

