## Supplementary figures for "Pan-cancer analysis of the ion permeome reveals functional regulators of glioblastoma aggression"

### Supplementary information

### Bahcheli, Min *et al.* (2023)

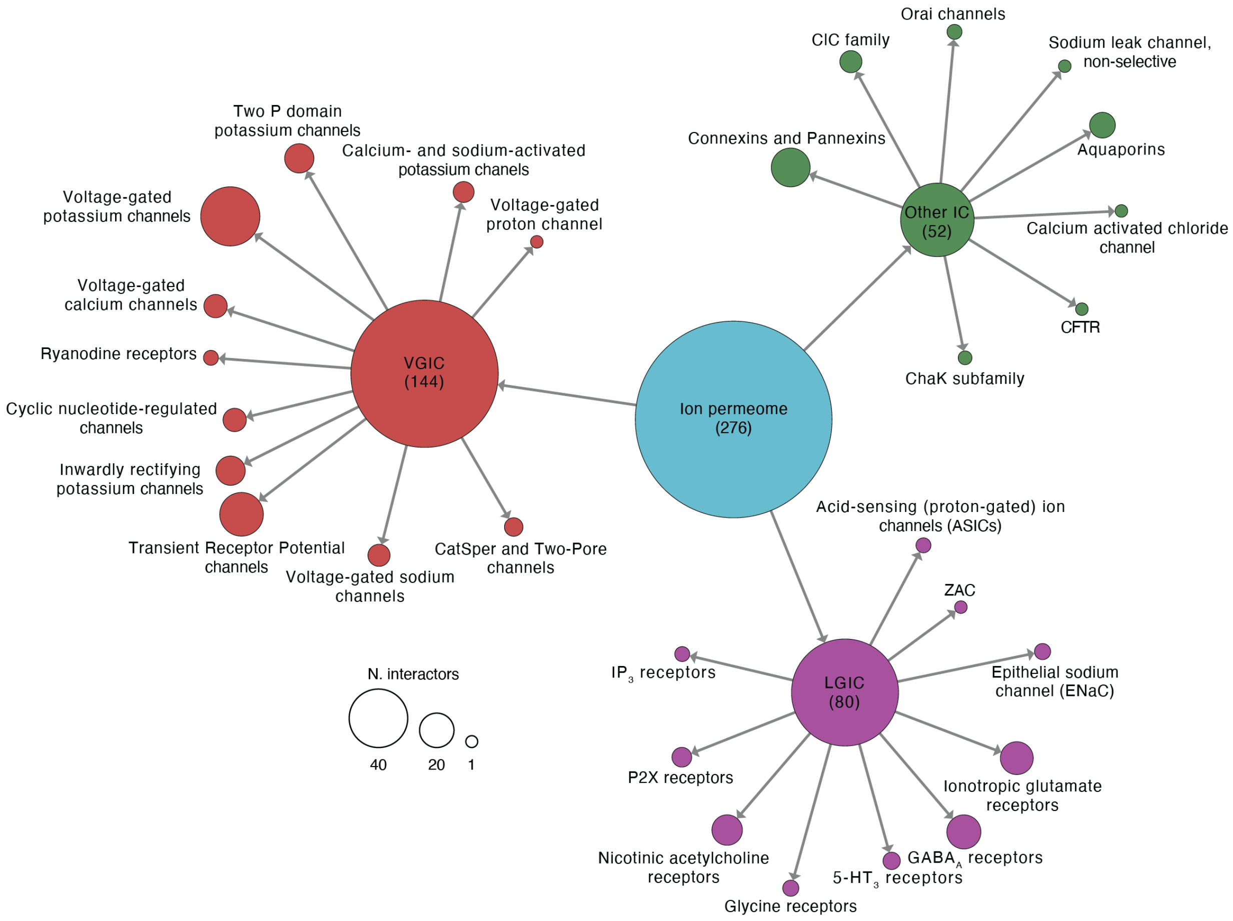


**Figure S1. Classification of ion permeome (IP) proteins used in our study.** Genes encoding IP proteins with known specific inhibitors were obtained from the Guide to Pharmacology database and filtered to include genes with expression profiles in TCGA. The network shows a hierarchical classification of IP proteins by type or family (nodes) with arrows indicating subclasses derived from larger classes. Bubble size reflects the number of genes within each type or family. Node size reflects the number of genes within each family. Node color reflects major IP gene classes. Other ion channels (IC), voltage-gated ICs (VGIC), and ligand-gated ICs (LGICs) represent the major classes.

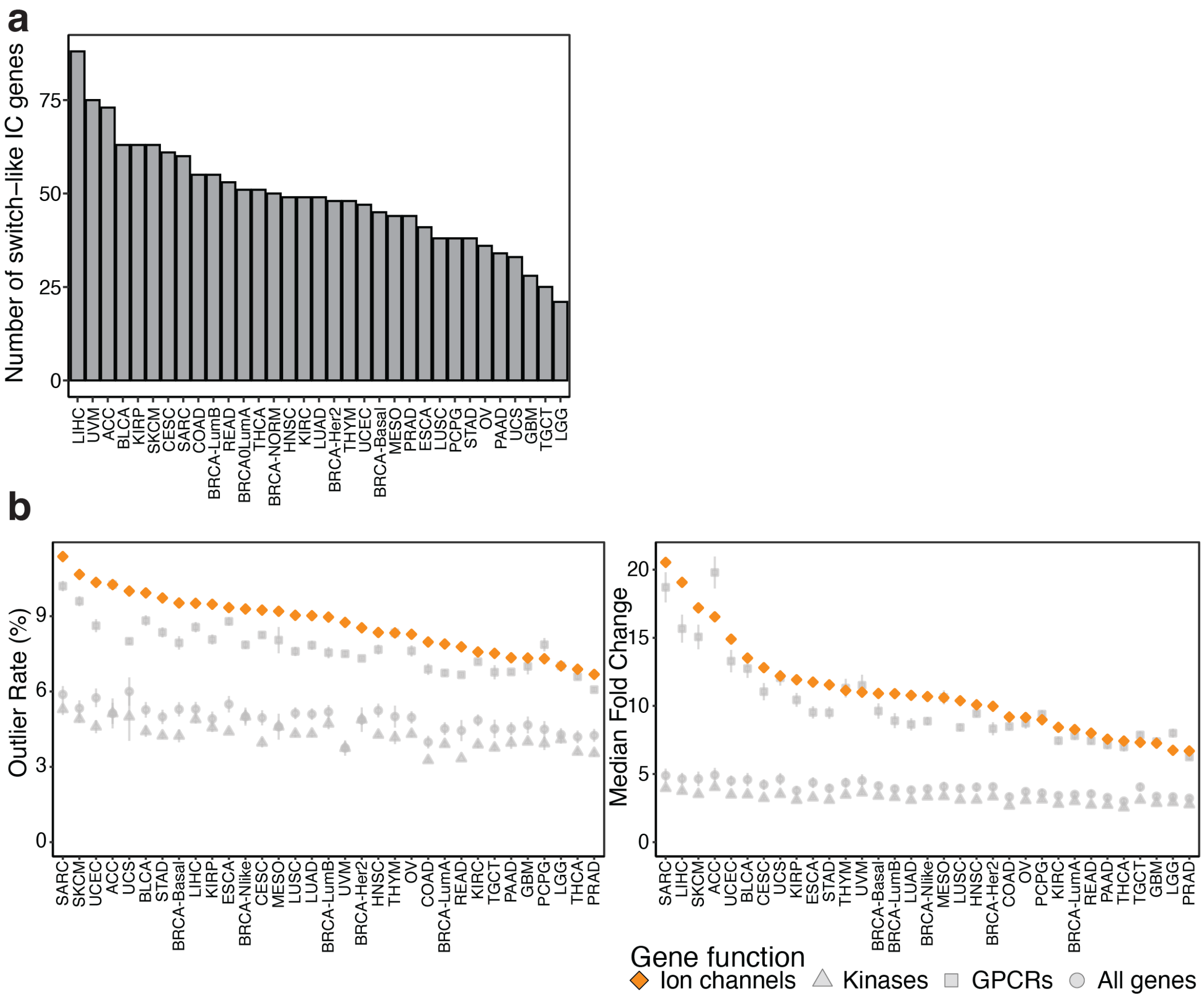


**Figure S2. Significantly elevated ion expression of IP genes in subsets of cancer samples. (a)** Statistical analysis of elevated IP gene expression using permutation tests in individual cancer types of the TCGA cohort. ICs (shown in orange) show higher fractions of cancer samples with highly elevated expression (left) and higher increase in expression relative to other samples of the same cancer type (right). Control genes (gray) include two major classes of drug targets, kinases and GPCRs (triangles and squares, respectively), and all protein-coding genes (circles). These control gene sets were sampled randomly 10,000 times. Sets of 276 genes were sampled as controls to match the number of IP genes we analysed. Error bars show one standard deviation. **(b)** The number of IP genes with switch-like expression patterns in each cancer type. Bar plot shows the number of genes for which the median expression in given cancer type was zero, while a minority of samples had elevated expression of the gene.

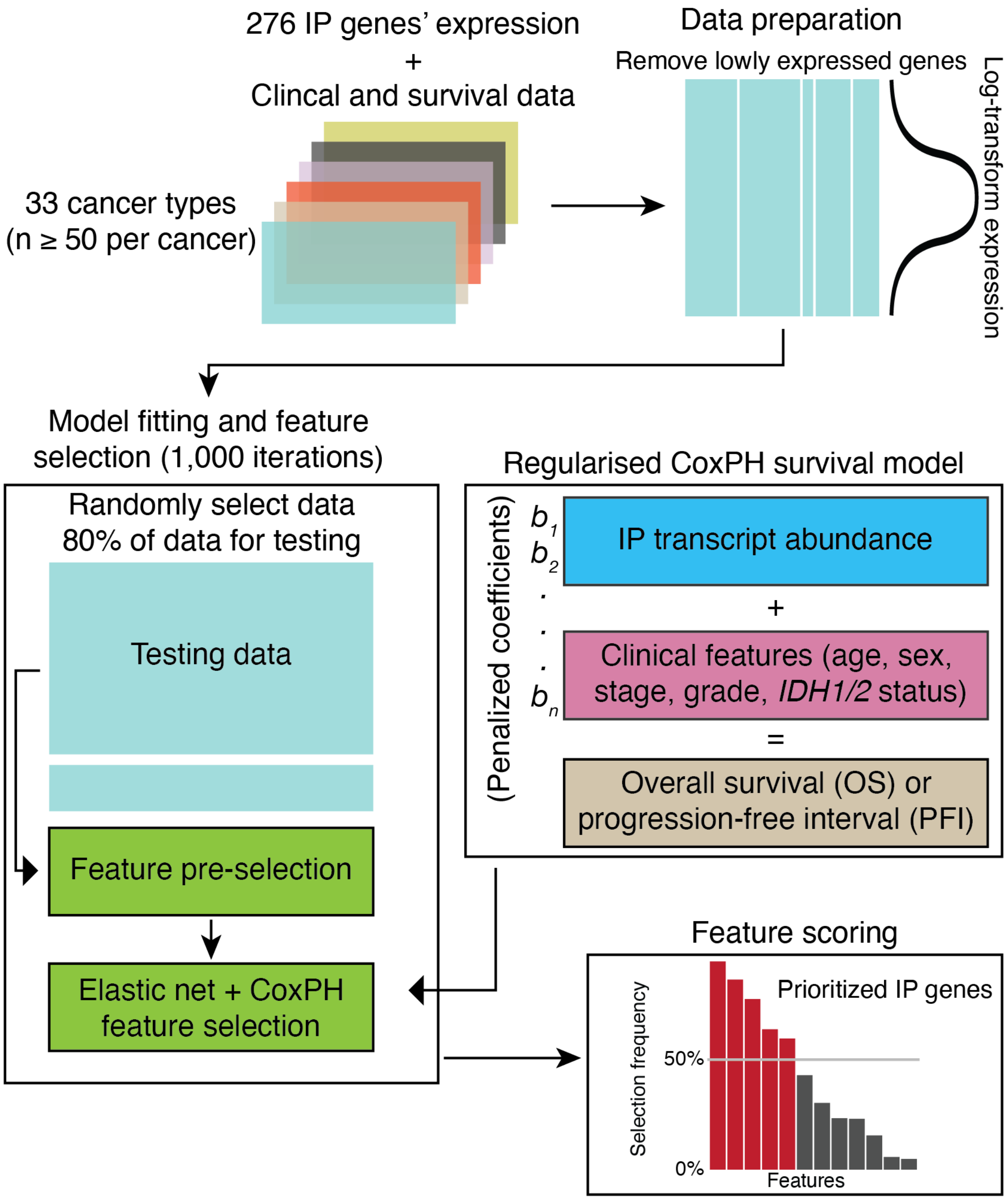


**Figure S3. Overview of the machine learning pipeline for identifying survival-associated ion permeome (IP) genes.** IP gene expression was evaluated for significant survival associations individually for 33 different cancer types of The Cancer Genome Atlas (TCGA). IP gene expression was log-normalized and used as a predictive feature in a machine learning framework. Regularised Cox proportional hazard (CoxPH) models were trained on iterations of 80% of samples within each cancer type, using patient IP gene expression and clinical factors (patient age and sex, tumor stage, grade, and *IDH1/2* mutation status) as predictive variables and patient survival as the response variable. Within each iteration, features were first pre-selected using univariate CoxPH models trained, and only genes significantly associated with patient survival ($P < 0.1$, Wald test) were selected for the multivariate model. After 1,000 iterations of machine learning feature prioritisation, IP genes associated with patient survival in at least half (>50%) of models were selected for further interpretation.

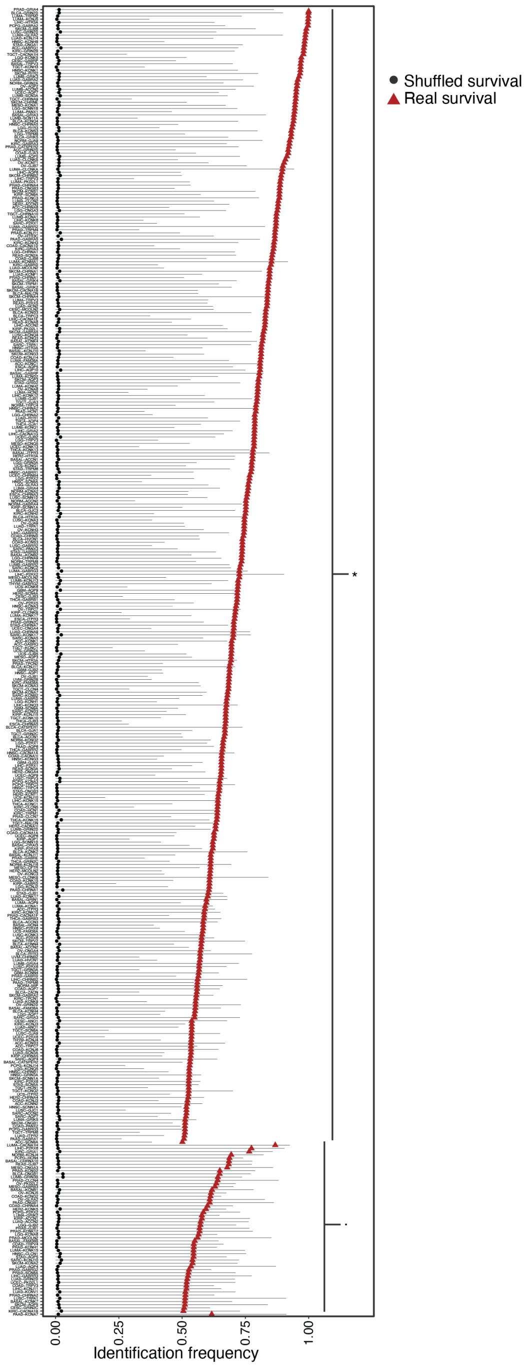


**Figure S4. Machine learning identifies IP genes associated with OS in GBM.** Elastic net feature selection identifies IP genes associated with patient survival in true datasets significantly more frequently than expected from permuted survival datasets. The analysis for IP gene selection was repeated on randomly shuffled patient survival data to discover survival-associated IP genes. Survival data was randomly permuted among patients over a series of 100 iterations. The frequencies of IP genes identified in true data (red dots) were compared to 100 iterations of shuffled data (black dots with 95% confidence intervals). Empirical *P*-values are shown (• $P < 0.1$, * $P < 0.05$).

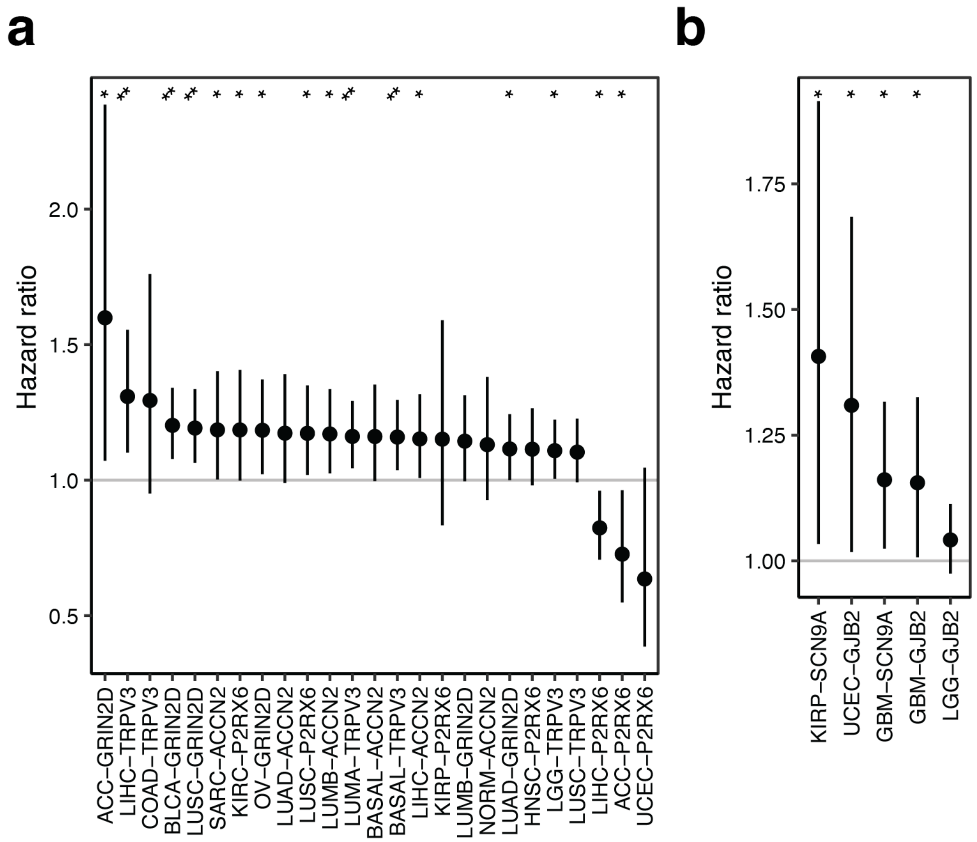


**Figure S5. Prioritized IP gene expression associates with patient survival.** Multivariate hazard ratios (HR) of IP genes prioritized in other cancer types (a) and in GBM (b). CoxPH survival models were trained on single IP gene expression and common clinical variables (age, sex, tumor stage and grade, and *IDH1/2* mutations). Median multivariate HR is shown with 95% confidence intervals. Wald test P-values are shown (* $P < 0.05$, ** $P < 0.01$).

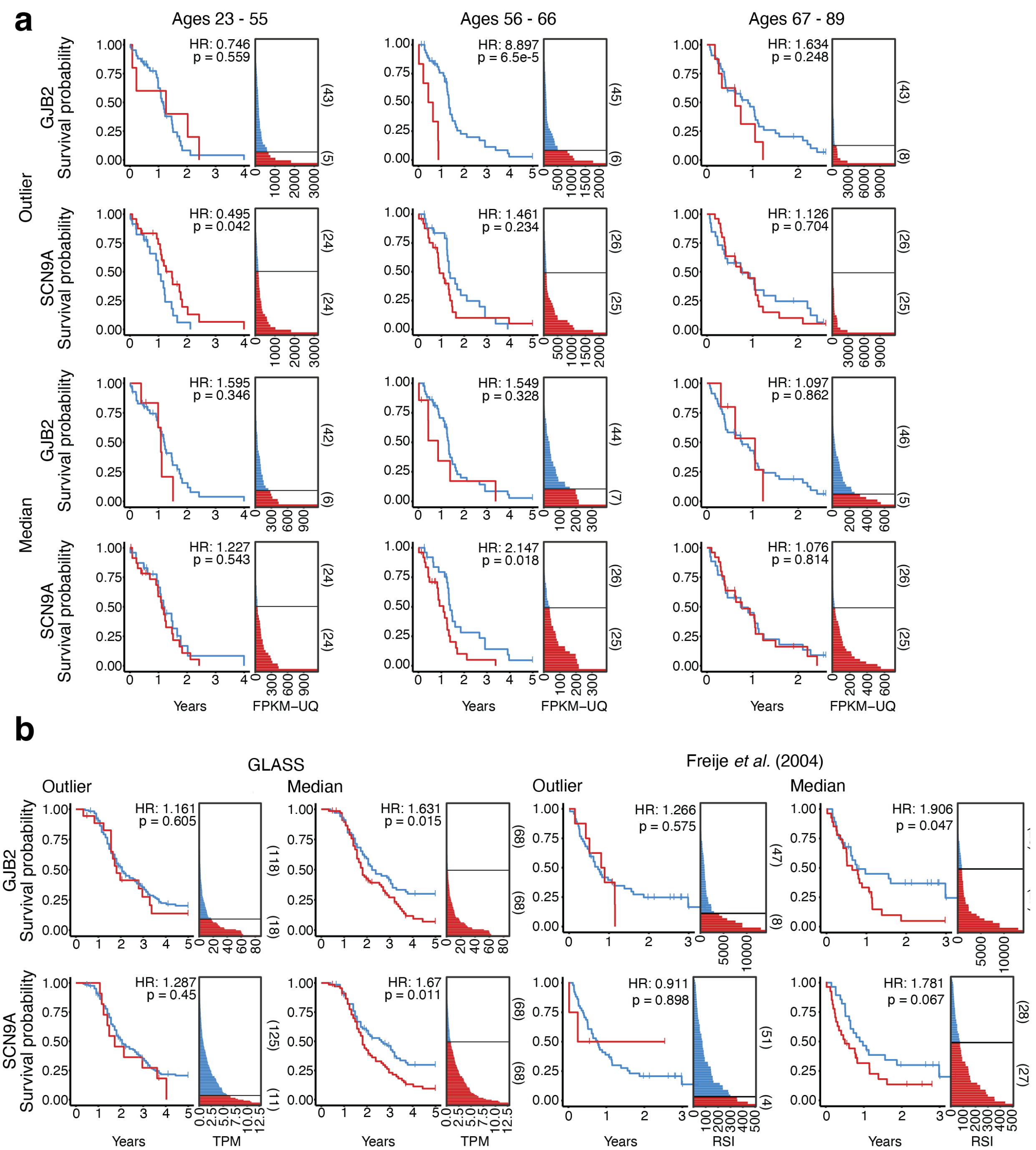


**Figure S7. *GJB2* and *SCN9A* expression is associated with patient overall survival (OS) in GBM.** Kaplan-Meier curves for GBM patient OS in TCGA **(a)** and two independent cohorts **(b)**. Samples were split into high or low risk groups by their expression levels by either a median or outlier cut-off. For each plot, the Kaplan-Meier survival curves (left) and sample expression (right) is shown, along with the number of samples classified into each group, and the hazard ratio (HR) and P-value (p) for each univariate CoxPH survival model trained on risk group classifications. For the TCGA cohort, patients were trichotomized by age into three groups of equal size.

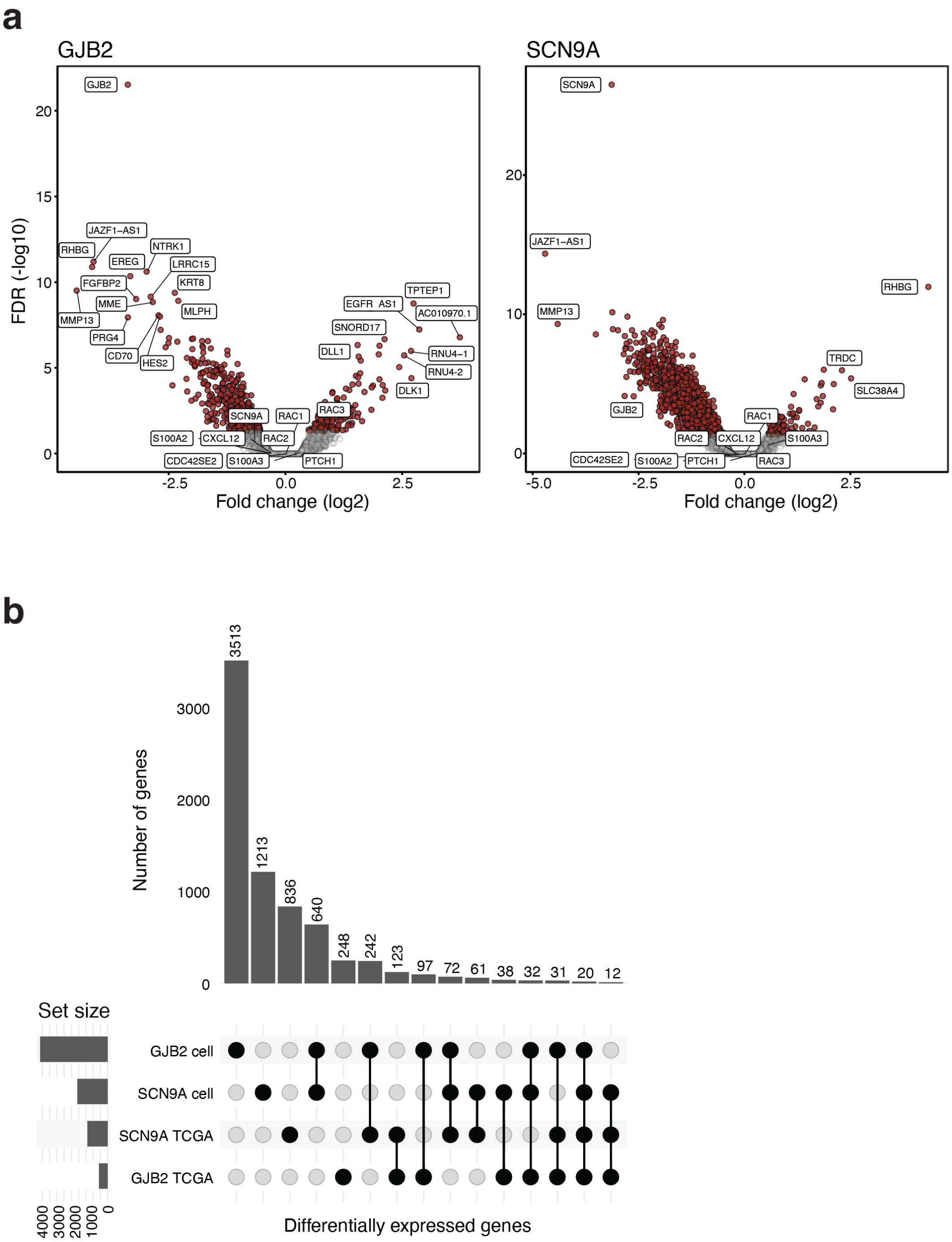


**Figure S7. Differential gene expression analysis (DGEA) results of samples with high versus low expression of *GJB2* and *SCN9A* in TCGA. (a)** Volcano plot of differentially expressed genes (DEGs) observed in TCGA GBM samples dichotomized by median expression of *GJB2* (left) and *SCN9A* (right). Significant DEGs are shown in red (EdgeR; FDR < 0.05) and genes associated with tunnelling nanotubes or with the most significant changes in expression are labelled. **(b)** Comparison DEGs from TCGA (panel A) with genes identified in *GJB2* or *SCN9A* knockdown experiments in patient-derived GBM cell lines. Bars show the number of genes found in the analyses of cell line experiments or patient GBMs in TCGA. Dot-and-lines visualisations show the combinations of DGEAs. The number of genes identified in each DGEA (set size) is shown as horizontal bars.
